# CIRCA learns identifiable cell- and niche-level programs and their interactions with tissue pathology from spatial transcriptomics

**DOI:** 10.64898/2026.11.17.767388

**Authors:** Tyler C Lovelace, Chia-Hao Tung, Nitesh Enduru, Zhongming Zhao

**Affiliations:** Center for Precision Health, McWilliams School of Biomedical Informatics, The University of Texas Health Science Center at Houston, Houston, TX 77030, USA; Department of Epidemiology, Human Genetics and Environmental Sciences, School of Public Health, The University of Texas Health Science Center at Houston, Houston, TX 77030, USA; Faillace Department of Psychiatry and Behavioral Sciences, McGovern Medical School, The University of Texas Health Science Center at Houston, Houston, TX 77030, USA

## Abstract

Interactions among cells and their local microenvironments shape tissue organization and function, while their dysregulation can contribute to disease. In Alzheimer’s disease (AD), the complex responses of glial cells and neurons to amyloid-β pathology can both mitigate and exacerbate neuronal injury. Spatial transcriptomics enables investigation of these relationships *in situ*, but recovering interpretable transcriptional programs and their interactions from spatial transcriptomics remains challenging, in part because conventional nonlinear models are non-identifiable. Here we present CIRCA (Causal Interaction Representation for Cellular Architectures), a self-supervised causal representation learning framework that jointly learns identifiable cell- and neighborhood-level transcriptional programs and the directed interactions among them under explicit assumptions. It can also be extended to incorporate additional modalities such as pathology. Benchmarked against five existing methods across seven cell-resolution datasets, CIRCA achieved the highest biological accuracy in six and the highest overall score in all five multi-section datasets, while recovering higher-order anatomical organization. In an AD mouse model, incorporating amyloid-β pathology revealed four region-specific plaque niches enriched for microglia. Plaque-associated microglia shared a core disease-associated response whose strength varied by region, and AD genetic risk was preferentially enriched in hippocampal and cortical plaque-associated microglia. Reproducible latent factors linked to cell-level amyloid burden separated phagocytic from inflammatory microglial programs. Applied to a human colorectal cancer dataset, CIRCA resolved patient-specific tumor niches, conserved stromal remodeling niches, and distinct macrophage states, highlighting its versatility. Together, these results establish CIRCA as a methodological advance over existing approaches, enabling deeper insights into tissue organization and the molecular dependencies linking cells to their local environments.

## Introduction

Multicellular tissues exhibit complex biological functions that emerge from interactions among individual cells and their local microenvironments (1). These interactions contribute to the formation and maintenance of tissue architecture and distinct cellular niches, characterized by their cellular composition, molecular states, and spatial organization (2, 3). Disruption of cellular states or their microenvironments can alter tissue organization and contribute to disease (4). For example, in Alzheimer’s disease (AD), extracellular amyloid-β (Aβ) plaques induce spatially organized glial and neuronal responses, whose interactions with local pathology can both mitigate and exacerbate amyloid accumulation, neuroinflammation, and neuronal injury (5–8). Understanding how these interactions reshape tissue function requires approaches that jointly characterize cellular states and their relationships with the local microenvironment. Recent advances in cell-resolution spatial transcriptomics technologies (9–17) provide an opportunity to systematically investigate these relationships directly within intact tissues (4, 18).

Many existing computational methods identify spatial domains and cellular niches by integrating gene expression with spatial proximity (19–30), while others model how local microenvironments influence cellular gene expression or infer spatially resolved cell-cell communication (31–34). However, few approaches jointly learn the underlying transcriptional programs and their interactions. NicheCompass addresses this challenge by incorporating prior knowledge from pathway and cell-cell communication databases to learn interpretable spatial gene programs (35). However, reliance on curated knowledge can limit the discovery of uncharacterized interactions, while its autoencoder-derived *de novo* programs are not guaranteed to recover a unique representation of the underlying biology. Thus, jointly recovering reproducible, interpretable transcriptional programs and their interactions without requiring predefined biological knowledge remains an important challenge.

This challenge reflects a fundamental limitation of conventional nonlinear representation learning approaches, including autoencoders: non-identifiability. A model is said to be non-identifiable if multiple distinct latent representations can be learned to explain the same observations, even with unlimited data (36–40). Causal representation learning seeks to overcome this limitation by introducing explicit assumptions about the underlying latent variables and their causal relationships (41–44). Rather than directly modeling relationships between observed features (*e.g.* gene expression), these approaches seek to identify the latent factors (*e.g.* transcription factor activity) responsible for generating observations and the cause-effect interactions among them. Recent theoretical advances have established conditions under which both latent factors and their interactions can be identified when observations are partitioned into groups containing distinct but interacting latent variables (42). Spatial transcriptomics provides a natural setting for this framework, as individual cells and their local neighborhoods contain distinct but interacting transcriptional programs. Moreover, because higher-order tissue organization can emerge from local cellular interactions (1), modeling these dependencies may capture broader anatomical structure without explicitly representing global spatial relationships.

Here, we introduce CIRCA (Causal Interaction Representation for Cellular Architectures), a self-supervised framework that applies grouped causal representation learning to spatial transcriptomics. CIRCA jointly learns identifiable cell-intrinsic and neighborhood-level transcriptional programs together with a sparse, directed interaction structure linking these programs under explicit assumptions about the data-generating process. Across seven spatial transcriptomics datasets, CIRCA achieves strong biological accuracy, spatial continuity, and multi-section integration while recovering higher-order anatomical organization from local cell-neighborhood pairs. We further extend CIRCA to incorporate Aβ pathology measurements as an additional group in an AD mouse model, resolving brain region-specific plaque microenvironments and interpretable transcriptional programs linked to pathology. To demonstrate its broad utility, we applied CIRCA to human colorectal cancer spatial transcriptomics data, revealing patient-specific malignant niches alongside shared patterns of tumor-associated stromal remodeling. Together, these results establish CIRCA as a framework for investigating spatial tissue organization and the molecular dependencies linking cells to their local environments.

## Results

### CIRCA learns identifiable cell–microenvironment representations

CIRCA employs causal representation learning (42) to identify latent cell- and niche-level transcriptional programs and their directed interactions. From cell-resolution spatial transcriptomics, it constructs a spatial k-nearest neighbor graph and represents each cell and its local neighborhood as separate feature groups, which form a cell-neighborhood pair (**Fig. 1a, step 1**). A shared multilayer perceptron (MLP) backbone followed by group-specific encoders maps each group’s expression profiles to cell- and niche-level latent factors (**Fig. 1a, step 2**). During training, data augmentation improves robustness to technical variation.

**Figure 1.**
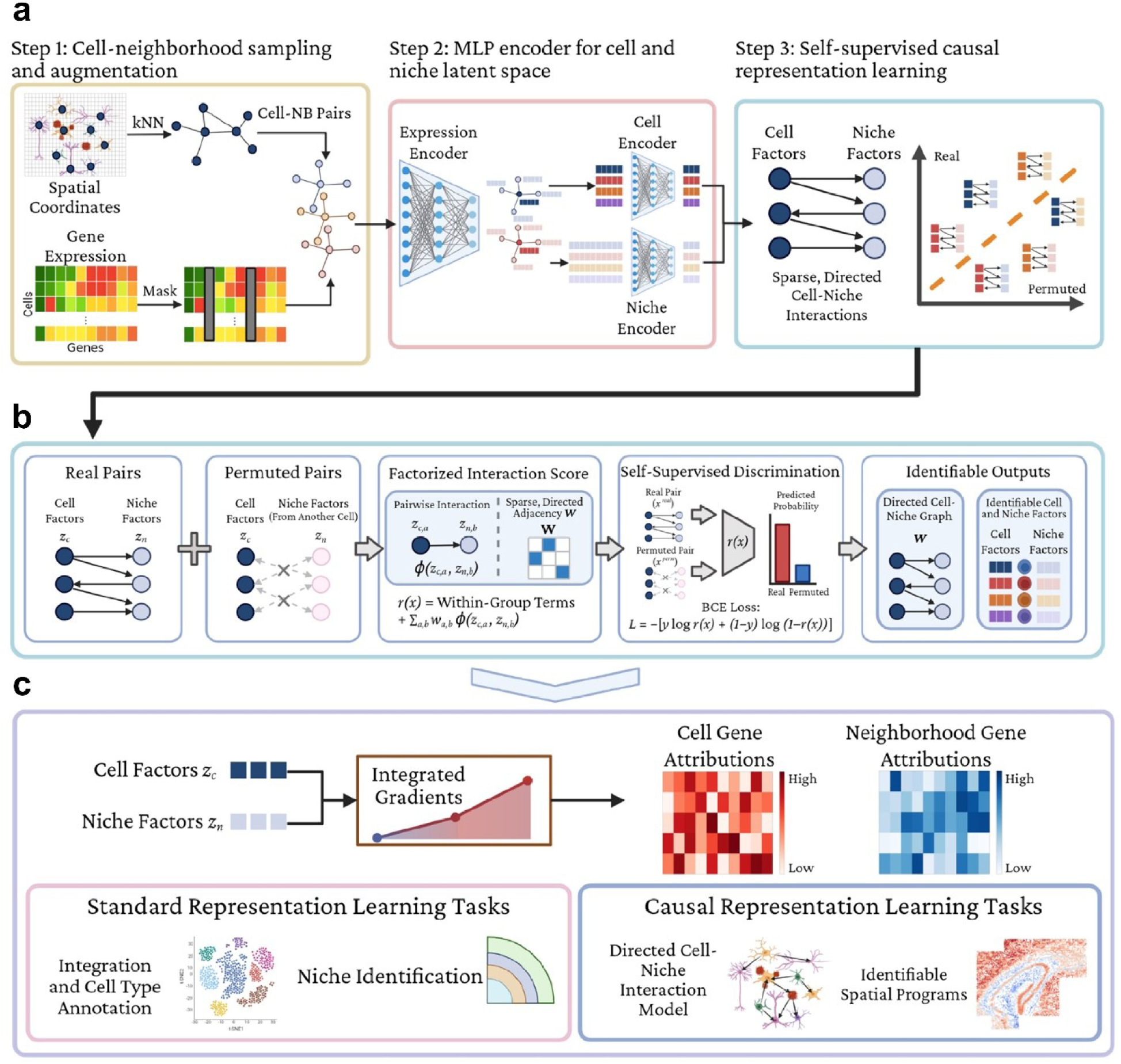
CIRCA: Causal Interaction Representation for Cellular Architectures. a) Model overview. CIRCA takes cell-resolution spatial transcriptomics data as input. **Step 1:** Slide-specific k-nearest neighbor graphs are built from spatial coordinates. Each cell and its neighborhood are treated as separate feature groups and sampled as cell–neighborhood pairs, with data augmentation to reduce sensitivity to technical variation. **Step 2:** A shared multilayer perceptron (MLP) backbone embeds the cell and neighbor expression profiles, which are then split and encoded into cell- and niche-level latent spaces. **Step 3:** A self-supervised causal representation learning objective recovers the latent factors and their directed interactions by discriminating real from permuted cell–neighborhood pairs. **b) Self-supervised causal representation learning.** Permutation preserves each group’s marginal distribution but breaks the dependencies between groups introduced by causal interactions. The discriminator uses a factorized interaction score weighted by a sparse, directed adjacency matrix *W*, constraining it to follow the assumed causal data-generating process. Under the model’s assumptions, optimal discrimination with binary cross-entropy (BCE) loss therefore requires recovering the latent factors and their directed cross-group interactions. **c) Biological interpretation.** Integrated Gradients attribute cell- and niche-level latent factors to genes and pathways, supporting standard spatial transcriptomics analyses as well as the recovery of reproducible spatial transcriptional programs and their interactions.

CIRCA jointly learns these latent representations and their sparse, directed interactions using a self-supervised objective (**Fig. 1a, step 3; Fig. 1b**). Independently permuting cells and neighborhoods within each tissue section yields negative pairs that preserve each group’s marginal distribution but break the causal relationships between groups. A discriminator distinguishes observed from permuted pairs using a factorized interaction score, in which pairwise latent interactions are modeled by a shared asymmetric nonlinear function and weighted by a sparse, directed adjacency matrix. Under the assumptions about the data-generating process described in **Methods**, this formulation recovers the latent factors up to permutation and component-wise invertible transformations, and the interaction structure up to permutation, scaling, and matrix transposition. Because negatives are drawn from within tissue sections, CIRCA supports multi-slide integration without an explicit batch-correction penalty. CIRCA’s grouped framework can also incorporate additional modalities, such as pathological measurements from paired immunofluorescence imaging, as additional groups.

Beyond conventional tasks such as cell typing, niche identification, and integration, CIRCA’s identifiable latent representations recover reproducible, interpretable transcriptional programs and their interactions without employing predefined gene sets or interaction databases (**Fig. 1c**). Integrated Gradients (IG) (45) then quantifies gene-level contributions to individual latent factors, enabling gene- and pathway-level interpretation of how cells respond to their local microenvironment.

### CIRCA recovers spatial domains and tissue architecture

We benchmarked CIRCA against five spatial transcriptomics methods across seven datasets spanning multiple technologies and gene panels ranging from 33 to 6,000 genes (**Table 1**). CIRCA achieved the highest biological accuracy in six of seven datasets and ranked second in the remaining dataset (**Fig. 2a, S1**). CIRCA also performed consistently well in spatial continuity and slide integration, achieving the highest overall score in all five multi-section datasets and ranking second to BANKSY in the two single-section datasets. When aggregated across benchmark datasets, CIRCA significantly outperformed each competing method in overall performance (p<0.05).

**Figure 2.**
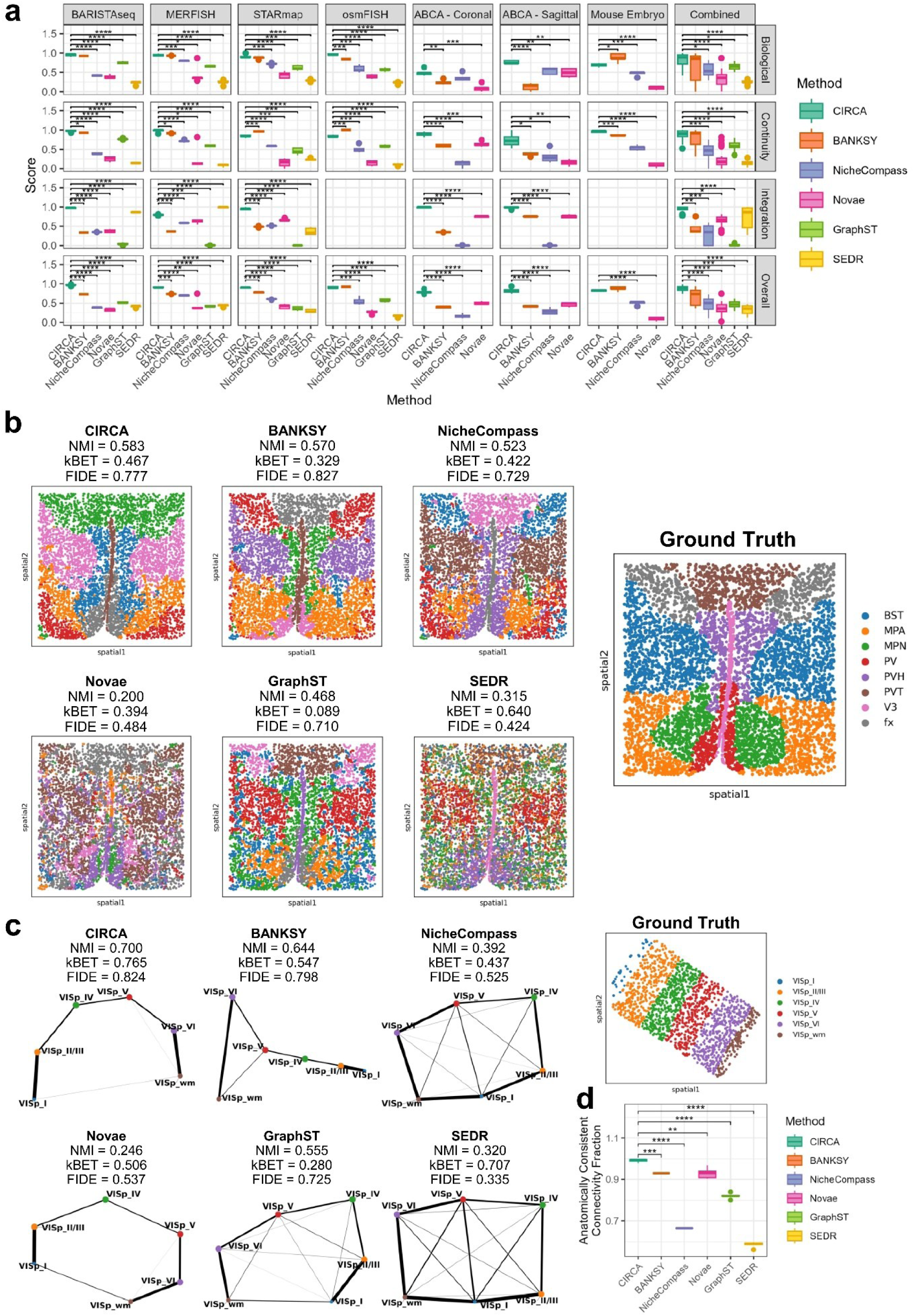
CIRCA improves niche identification, multi-slide integration, and recovery of tissue architecture. **a)** Benchmarking results for CIRCA and five competing methods across seven annotated datasets on biological accuracy, spatial continuity, multi-slide integration, and overall performance. Boxplots depict performance across 5 random seeds; the center line denotes the median, box limits denote the upper and lower quartiles, and whiskers denote 1.5x the interquartile range; Welch’s t-test (within dataset) and mixed effects model (combined) Holm adjusted p-values: * p<0.05, ** p<0.01, *** p<0.001, **** p<0.0001. **b)** Ground truth annotations and unsupervised clusters from each method’s median-NMI run on a single slide of the MERFISH hypothalamus dataset, with NMI, FIDE, and kBET values for the displayed runs. **c)** Ground truth annotations for the BaristaSeq dataset and PAGA graphs computed on each method’s median-NMI embedding, with the same metrics shown. **d)** Fraction of PAGA connectivity that is anatomically consistent for each method across five seeds. Boxplots and p-values are the same as in **a**.

**Table 1:** Characteristics of spatial transcriptomics datasets used in this work. Cell and gene numbers are after preprocessing and quality control. The number of domains refers to the number of reference domain labels in benchmarking datasets.

| Dataset | Technology | # Sections | # Cells | # Genes | # Domains | Reference |
|---|---|---|---|---|---|---|
| **BaristaSeq** | BaristaSeq | 3 | 4,754 | 77 | 6 | (11) |
| **MERFISH** | MERFISH | 5 | 26,901 | 155 | 8 | (46) |
| **osmFISH** | osmFISH | 1 | 4,839 | 33 | 11 | (12) |
| **STARmap** | STARmap | 3 | 3,190 | 166 | 4 | (88) |
| **ABCA – Coronal** | MERFISH | 8 | 239,857 | 1,122 | 18 | (89) |
| **ABCA – Sagittal** | MERFISH | 3 | 239,795 | 1,122 | 23 | (89) |
| **Mouse Embryo** | Stereo-seq | 1 | 281,377 | 6,000 | 22 | (14) |
| **AD Mouse Model** | CosMx | 4 | 36,455 | 900 | N/A | (49) |
| **Colorectal Cancer** | Visium HD | 5 | 937,559 | 9,472 | N/A | (16) |

The MERFISH mouse hypothalamus dataset (46) illustrates CIRCA’s ability to recover spatially contiguous niches that align with known anatomical organization across consecutive tissue sections (**Fig. 2b**). CIRCA achieved the highest median normalized mutual information (NMI; 0.583), with BANKSY (0.570) and NicheCompass (0.523) ranking second and third, respectively, while GraphST (0.468), SEDR (0.315), and Novae (0.200) showed weaker correspondence with the annotated regions. These spatial maps from the median-performing runs for each method illustrate CIRCA’s ability to recover spatially and anatomically coherent domains (**Fig. 2b, left**).

Beyond domain identification, we evaluated whether the learned niche representations preserved higher-order anatomical relationships using three BaristaSeq sections from the mouse visual cortex (11). Partition-based graph abstraction (PAGA) graphs (47) constructed from the learned representations and published anatomical annotations revealed that CIRCA most closely reproduced the expected laminar organization, progressing from cortical layer I through layers II/III, IV, V, and VI to the white matter (**Fig. 2c**). Quantification of the fraction of PAGA connectivity consistent with anatomical adjacency confirmed that CIRCA concentrated the greatest proportion of PAGA connectivity on anatomically adjacent layers, with fewer spurious connections between distant regions (p<0.05; **Fig. 2d**). Together, these results demonstrate that CIRCA preserves both spatial domain identity and the broader anatomical relationships among tissue compartments.

### CIRCA resolved region-specific Aβ-associated niches while preserving anatomical tissue organization

In Alzheimer’s disease, amyloid-β (Aβ) is linked to AD progression and symptom severity (48). To investigate how Aβ pathology reshapes the brain microenvironment in AD, we applied CIRCA to a publicly available CosMx 1K dataset generated from hippocampal sections of four 18-month-old AppNL-G-F knock-in mice (49), the only publicly available cell-resolution spatial dataset with matched Aβ measurement. Paired Aβ immunofluorescence (IF) measurements were incorporated as an additional feature group containing the mean and maximum Aβ IF intensities of each cell and its local neighborhood. This enabled CIRCA to jointly model cell transcriptional states, neighborhood-associated features, pathological measurements, and their interactions within a common analytical framework (**Fig 3a**).

**Figure 3.**
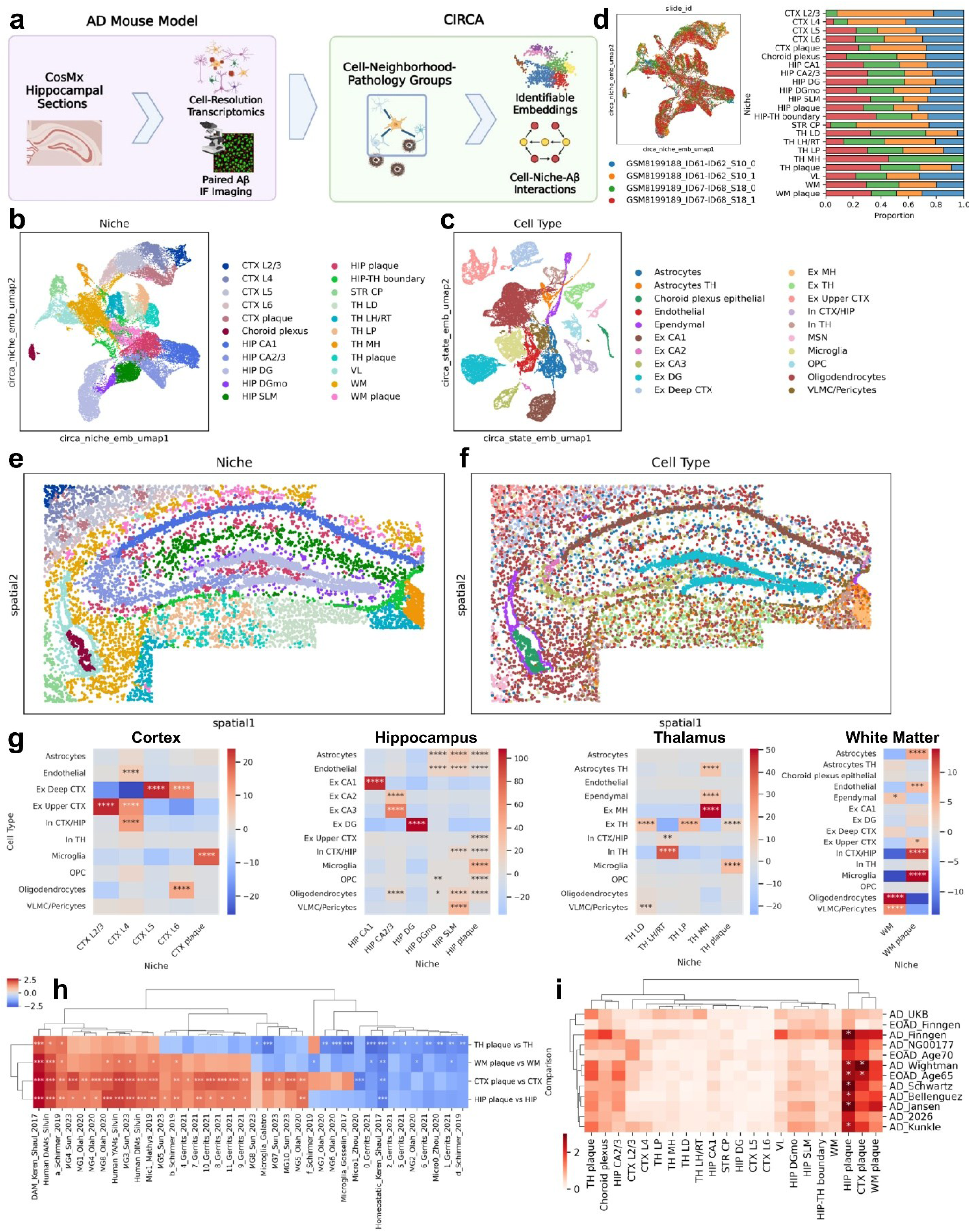
CIRCA resolved region-specific plaque niches and microglial responses in an AD mouse model. **a)** CIRCA is applied to a CosMx dataset of hippocampal sections from four humanized APP mice with paired Aβ immunofluorescence, jointly modeling cell, neighborhood, and pathology groups and their interactions. **b-c)** UMAPs of CIRCA niche (**b**) and cell-state (**c**) embeddings annotated with niche (**b**) and cell type (**c**) labels derived from Leiden clustering. **d)** Niche embedding UMAP colored by slide of origin, with the proportion of each niche contributed by each slide. **e-f)** Spatial distributions of CIRCA-identified niches (**e**) and cell types (**f**) in a representative hippocampal section. **g)** Adjusted standardized residuals from chi-square tests of cell type-niche association, stratified by brain region (CTX, HIP, TH, WM); Bonferroni-adjusted post hoc p-values: * p<0.05, ** p<0.01, *** p<0.001, **** p<0.0001. **h)** GSEA normalized enrichment scores of published microglial state signatures (50) in plaque-associated versus non-plaque microglia, shown for signatures significant in at least one region. False discovery rate (FDR): * <0.05, ** <0.01, *** <0.001. **i)** scDRS enrichment for GWAS-derived AD genetic risk across CIRCA niches. FDR: * <0.1, ** <0.05.

We first assessed whether CIRCA recovered established cellular identities and anatomical organization. Unsupervised clustering of CIRCA’s cell-state and niche embeddings recovered expected cell types and anatomical regions across the hippocampus and adjacent cortical, thalamic, white matter, and striatal regions (**Fig. 3b,c, S2**). The niche representations resolved distinct cortical layers (L2/3 through L6), hippocampal fields (CA1, CA2/3, and dentate gyrus), and thalamic nuclei (LD, LP, MH, and LH/RT) (**Fig 3e, S3**). Cell type clusters also followed expected anatomical distributions, with upper- and deep-layer excitatory neurons enriched in their corresponding cortical niches (p<0.0001; **Fig 3f,g**). Together, these results indicate that CIRCA captures complementary cellular and anatomical information across multiple spatial scales.

We next examined the consistency of anatomical representations across tissue sections. Shared anatomical niches were represented across tissue sections containing the corresponding structures, while anatomically restricted niches, including MH and cortical layers L2/3, were retained only in sections in which these structures were present (**Fig. 3d, S3**). These results indicate that CIRCA captures recurring anatomical organization while retaining section-associated variation in the analyzed tissue samples.

We subsequently examined how pathology-associated features were represented across anatomically distinct tissue environments. Notably, CIRCA identified four distinct Aβ plaque-associated niches in the cortex, hippocampus, white matter, and thalamus (**Fig 3b**). Each plaque-associated niche showed elevated mean and maximum Aβ IF intensities relative to surrounding tissue within the corresponding region (p < 0.0001; **Fig. S4a,b**). Despite differences in their regional cellular composition, all four plaque-associated niches were enriched for microglia relative to non-plaque niches within the corresponding brain region (p<0.0001; **Fig. 3g**). This observation is consistent with previous analyses of this dataset reporting heterogeneous cellular compositions and increased microglial accumulation in Aβ-associated niches (49). Importantly, CIRCA provides an identifiable framework in which these plaque-associated cellular environments can be examined alongside their anatomical and neighborhood-level molecular context.

### Plaque-associated microglia show regionally varying responses and enrichment for AD genetic risk

Microglia undergo substantial transcriptional changes in response to Aβ pathology, including the acquisition of disease-associated microglia (DAM) signatures (50). To characterize the regional variation of these responses, we performed pseudobulk differential expression analysis of microglia, aggregating transcript counts by tissue section and brain region (**Fig. S5**) and comparing plaque-associated niches with their corresponding non-plaque regions (**Fig. S6**). Microglia in cortical, hippocampal, and white matter plaque-associated niches shared a core pathology-associated transcriptional response, characterized by elevated expression of DAM genes (*Apoe*, *Cst7*, *Itgax, Spp1*) and reduced expression of homeostatic microglial markers (*P2ry12, Tmem119*; **Fig S6**) (51). Thalamic microglia showed similar but generally weaker changes, with no individual genes reaching significance. Consistent with regional variation in individual gene responses, correlations between gene-level fold changes were modest, with the highest concordance observed between the cortex and hippocampus (Spearman’s ρ = 0.45) and correlations of ρ ≤ 0.26 for other regional comparisons (**Fig. S6**).

We next assessed whether these regional responses were preserved at the level of established microglial state signatures. Gene set enrichment analysis using published microglial state signatures (50) revealed enrichment of the classical DAM signature (FDR < 0.001) and depletion of the homeostatic signature (FDR < 0.01) in all four regions (**Fig. 3h**). At the signature level, plaque-associated profiles were more concordant across regions (cortex-hippocampus ρ = 0.90; hippocampus-white matter ρ = 0.84). The thalamus remained less concordant with other regions (ρ = 0.52-0.69) and exhibited the weakest DAM enrichment. Thus, plaque-associated microglia retain a conserved DAM-like response across anatomically distinct regions, despite regional variation in response magnitude and gene-level composition, with the thalamus showing the most distinct profile.

To explore the genetic relevance of CIRCA-identified cell types and spatial niches to AD, we applied scDRS (52) to single-cell expression profiles and evaluated enrichment for genetic risk signals across these populations. Using scDRS, we computed cell-specific disease relevance scores derived from genome-wide association study (GWAS) summary statistics spanning nine late-onset AD (LOAD) (53–61) and three early-onset AD (EOAD) (59, 62) datasets (**Table S1**). Consistent with prior work (56, 63–65), microglia showed significant transcriptional enrichment for AD polygenic risk across the evaluated cohorts (5/9 LOAD; 1/3 EOAD; FDR < 0.1; **Fig. S7**). At the niche-level, significant genetic associations were predominately localized to hippocampal (6/9 LOAD; 1/3 EOAD) and cortical (1/9 LOAD; 1/3 EOAD) plaque-associated niches (**Fig. 3i**). When further stratifying major cell types by anatomical region, hippocampal plaque-associated microglia exhibited the most robust and consistent enrichment (8/9 LOAD, 2/3 EOAD). Cortical plaque-associated microglia also showed recurrent enrichment (6/9 LOAD, 2/3 EOAD), while fewer associations were observed for white matter and thalamus plaque-associated microglia (**Fig. S8**). Together, these results demonstrate that plaque-associated microglial populations, particularly within the hippocampus and cortex, are characterized by transcriptional enrichment of genes prioritized by AD GWAS. This finding underscores how local pathology and spatial microenvironments intersect to shape the genetic risk architecture of microglia in AD.

### CIRCA resolves interpretable factors and interactions with Aβ pathology

We next investigated the reproducibility and biological interpretability of CIRCA’s learned latent factors. To assess factor stability, we compared three independently trained models generated using different random seeds and train-validation splits, matching factor representations using held-out cell-neighborhood pairs. Factors with an absolute Spearman correlation greater than 0.6 were considered replicated. Among the 50 cell-level and 50 niche-level factors, 39 (78%) cell-level factors and 44 (88%) niche-level factors were replicated in at least two of three runs, with mean absolute correlations of 0.70 and 0.72, respectively, among replicated factors. Both Aβ pathology-associated factors were consistently recovered across all three runs (mean |ρ| = 0.98). These results demonstrate that most cell- and niche-level factors are reproducible across independent training runs, with particularly high stability for the pathology-associated factors.

The two pathology-associated factors captured distinct spatial scales of Aβ burden. The first, termed Cell Aβ Burden, correlated with cell-level Aβ immunofluorescence (IF) intensity (Spearman’s ρ = 0.67; **Fig. 4a**), while the second, Niche Aβ Burden, correlated strongly with mean neighborhood Aβ intensity (ρ = 0.91; **Fig. 4b**). Both factors distinguished plaque-associated niches from surrounding anatomical regions, with greater separation observed for Niche Aβ Burden (**Fig. S4c,d**). Thus, without supervision, CIRCA recovered distinct cellular and neighborhood-level representations of Aβ burden.

**Figure 4.**
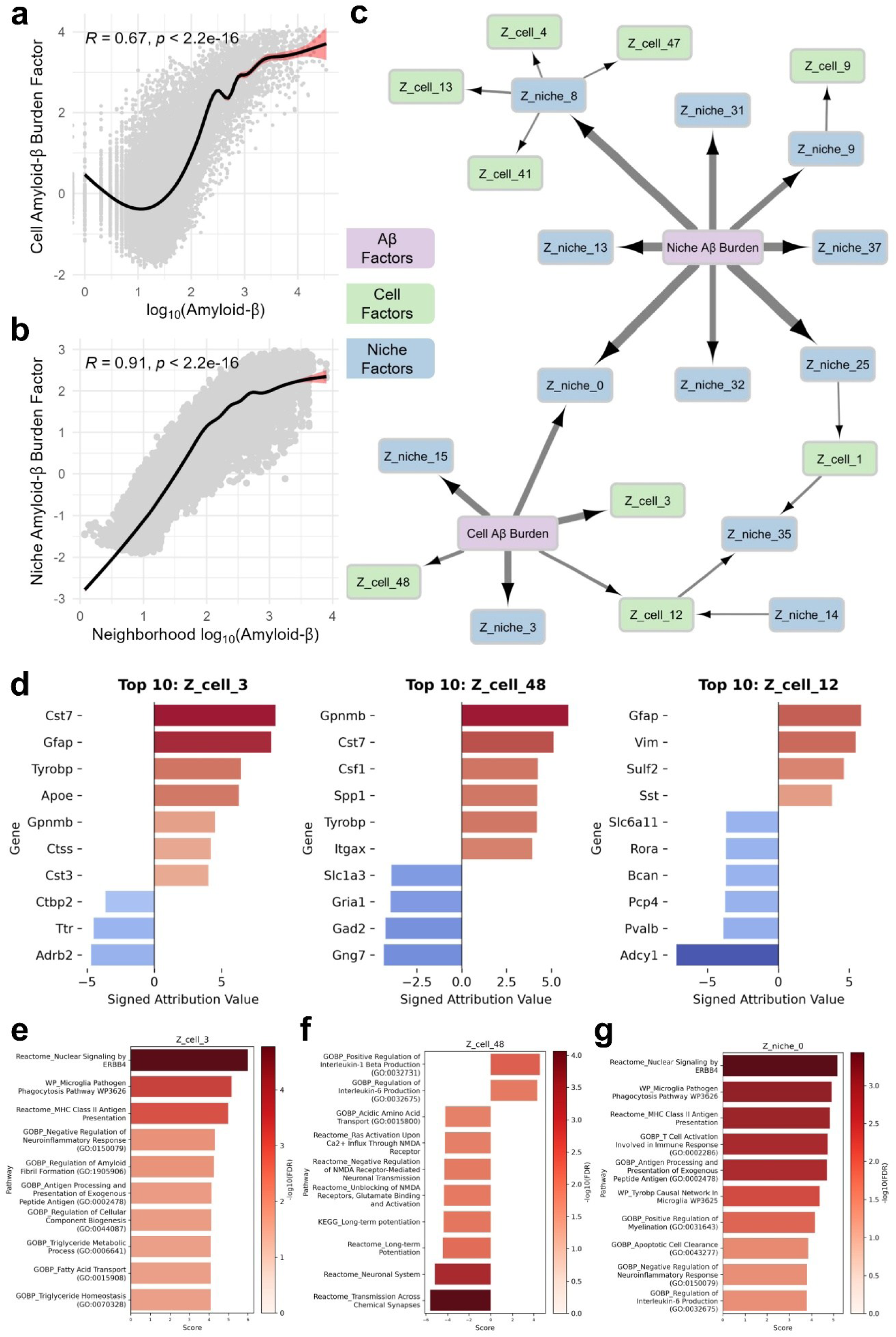
CIRCA captures cell- and niche-level transcriptional programs that interact with amyloid-β (Aβ) pathology. a-b) Cell Aβ Burden factor versus cell-level Aβ immunofluorescence intensity (**a**), and Niche Aβ Burden factor versus mean neighborhood Aβ intensity (**b**). Trendlines are estimated with a generalized additive model; Spearman correlation and unadjusted p-values are shown. **c)** Directed graphical model of interactions among replicated cell, niche, and amyloid latent factors, restricted to interactions elevated in plaque niches (FDR<0.01). For clarity, only factors within two edges of an amyloid pathology factor are shown. **d)** Genes with the ten largest absolute attributions for the three cell-level transcriptional programs 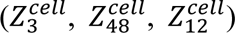 that interact directly with Cell Aβ Burden. Factors are oriented so that higher values correspond to higher Aβ burden; positive attributions indicate genes that increase the factor. **e-g)** The top ten pathways and biological processes with significant positive or negative enrichment (FDR < 0.05) in 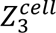 (**e**), 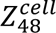 (**f**), and 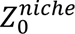 (**g**).

To investigate the transcriptional programs associated with these pathology factors in CIRCA’s interaction model, we used IG attributions (**Fig. 1c**) to interpret the gene-expression contributions to replicated cell- and niche-level factors associated with Aβ pathology. We focused on a subgraph comprising replicated latent factors within two edges of either Aβ pathology factor, retaining adjacencies whose interaction scores, or contributions to CIRCA’s discriminator logits, were significantly elevated in plaque-associated niches relative to other tissue regions (FDR < 0.01; **Fig. 4c**). Within this subgraph, Cell Aβ Burden was directly linked to three cell-level and three niche-level factors, including one niche factor also directly connected to Niche Aβ Burden. In contrast, Niche Aβ Burden was directly connected only to niche-level factors. This interaction structure revealed distinct patterns of model-inferred dependencies between cellular and neighborhood-level programs and the two pathology-associated factors, consistent with the different spatial scales they represent.

Gene-level attributions identified distinct cell-level transcriptional programs linked to Cell Aβ Burden (**Fig. 4d**). Two cell-level factors, 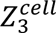 and 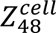 were characterized by the presence of DAM-associated genes (51), including *Cst7*, *Gpnmb*, *Spp1*, and *Tyrobp*. Notably, *Gpnmb* was a top contributor to both factors, ranking first in 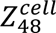 and fifth in 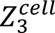. Supporting the relevance of *GPNMB* in human AD, a recent large-scale study identified *GPNMB*^+^ microglial populations associated with clinical AD dementia, cognitive decline, and neuropathological measures across diverse population groups and multiple brain regions (66). Another factor, 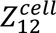, was associated with a disease-associated astrocyte (DAA)-like signature marked by *Gfap*, *Sulf2*, and *Vim* (67). Several Aβ-associated niche factors showed overlapping gene-level signatures (**Fig. S9**), characterized by positive contributions from DAM-associated genes and negative contributions from homeostatic microglial markers such as *Tmem119* (51). These overlapping signatures made functional distinctions among individual niche factors less apparent from their leading gene-level attributions alone.

Pathway-level attribution further revealed the functional distinctions among these factors (**Fig. 4e-g**). At the cell level, one DAM-associated factor, 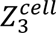, was enriched for pathways related to microglial phagocytosis, fatty-acid metabolism, and MHC class II antigen-presentation pathways (FDR < 0.05; **Fig. 4e**). In contrast, 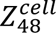, captured an axis contrasting inflammatory microglial signaling programs in regions with elevated cell-level Aβ burden with neuronal signaling programs elsewhere (FDR < 0.05; **Fig. 4f**). At the niche level, 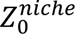, which was directly connected to both pathology factors, was enriched for antigen presentation, microglial phagocytosis, apoptotic cell clearance, neuroinflammatory regulation, and ERBB4-related signaling pathways (FDR < 0.05; **Fig. 4g**). By comparison, 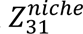, which was directly linked only to Niche Aβ Burden, shared neuroinflammatory and phagocytic signatures but additionally exhibited enrichment for TYROBP signaling, regulation of IL-6 production, and immune cell activation (FDR < 0.05; **Table S2**). Together, these results demonstrate that CIRCA resolves distinct transcriptional programs and their inferred interactions with Aβ pathology, even when their leading gene-level attributions are similar.

### CIRCA resolves patient-specific tumor niches and shared stromal remodeling in colorectal cancer

To evaluate CIRCA across disease settings and at a transcriptome-wide scale, we applied it to a Visium HD dataset comprising three human colorectal cancer (CRC) sections and two normal adjacent tissue (NAT) sections, containing more than 900,000 cells and 9,000 genes after cell segmentation and quality control (**Fig. 5a**). CIRCA learned integrated cell- and niche-level representations across all five sections (**Fig. 5b,c**). In NAT sections, niche clustering recovered spatially organized colonic structures, including apical, middle, and deep mucosal niches, lamina propria, lymphoid follicles, submucosa, and muscular layers (**Fig. 5e, S10**), and PAGA recovered their expected connectivity (**Fig. 5d**). Notably, the niche representation distinguished the muscularis mucosae from the muscularis propria despite their transcriptional similarity, indicating that CIRCA captures anatomical distinctions that are not readily resolved by cell-state similarity alone.

**Figure 5.**
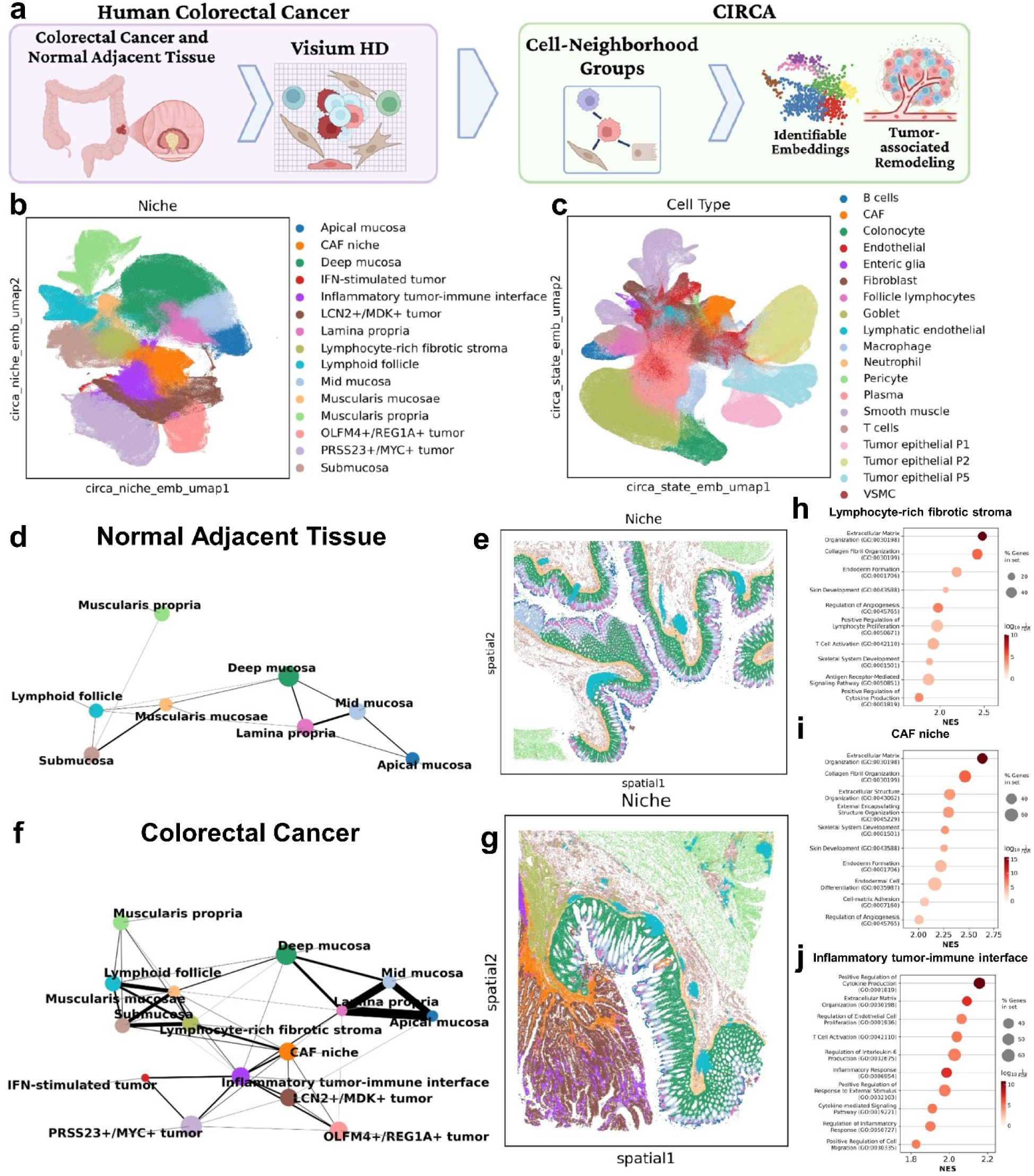
CIRCA captures shared patterns of stromal remodeling while preserving tumor heterogeneity in colorectal cancer. **a)** CIRCA is applied to a Visium HD dataset of three colorectal cancer (CRC) and two normal adjacent tissue (NAT) sections, jointly modeling cell and neighborhood groups and their interactions. **b-c)** UMAPs of CIRCA niche (**b**) and cell-state (**c**) embeddings annotated with niche (**b**) and cell type (**c**) labels derived from Leiden clustering. **d,f)** PAGA graphs of the spatial architecture of NAT (**d**) and CRC-remodeled tissue (**f**), inferred from CIRCA’s niche embeddings. **e,g)** Spatial distributions of CIRCA-identified niches in the P5_NAT (**e**) and P5_CRC (**g**) sections. **h-j)** The top ten Gene Ontology (GO) Biological Process (BP) terms with significant enrichment (FDR < 0.05) in the lymphocyte-rich fibrotic stroma (**h**), cancer-associated fibroblast (CAF) (**i**), and inflammatory tumor-immune interface (**j**) niches. NES: normalized enrichment score.

We next examined how CIRCA represented malignant tissue organization across the three CRC sections. In tumors, malignant cell heterogeneity poses a challenge for multi-sample integration because removing technical variation can also obscure biologically meaningful patient-specific differences (68). CIRCA preserved distinct patient-specific malignant niches, characterized by different transcriptional signatures: *OLFM4* and *REG1A* in P1, *PRSS23* and *MYC* in P2, and *LCN2* and *MDK* in P5, consistent with prior work showing malignant cells form stable, patient-specific transcriptional states (69) (**Fig. 5b**). P2 additionally contained an interferon-responsive tumor niche characterized by *CXCL10*, *ISG15*, *MX1*, *OAS2*, and *STAT1*. Compared to its primary tumor niche, this interferon-responsive niche exhibited higher macrophage and endothelial cell proportions (p < 0.0001; **Fig. S11**). These results indicate that CIRCA preserves patient-specific malignant transcriptional heterogeneity while resolving additional variation within individual tumors.

Despite this heterogeneity, CIRCA identified three tumor-associated stromal niches shared across CRC sections and absent from NAT (**Fig. 5f,g**). PAGA connectivity revealed these niches shared a consistent spatial organization from normal stroma toward tumor tissue. The first niche, termed lymphocyte-rich fibrotic stroma, was enriched for fibroblasts, plasma cells, and immune-related processes (**Fig. 5h, S11**). The second, a cancer-associated fibroblast (CAF) niche, showed enrichment for extracellular matrix (ECM) remodeling and angiogenesis (**Fig. 5i**). The third, an inflammatory tumor-immune interface, formed the principal boundary with malignant niches and was characterized by increased proportions of malignant epithelial and immune cells, together with enrichment for inflammatory and cytokine signaling pathways (**Fig. 5j, S11)**. Thus, CIRCA resolved shared tumor-associated stromal organization across the analyzed CRC sections while retaining patient-specific malignant heterogeneity.

### Distinct macrophage states occupy normal and tumor-associated niches

Macrophages were enriched in the CAF and tumor-immune interface niches, consistent with previous proximity-based analyses of macrophage populations in this dataset (16). To further characterize their transcriptional heterogeneity, we reclustered macrophages using CIRCA’s cell-state embeddings, identifying four distinct macrophage states (**Fig. S12a,b**). Two *SELENOP*+ states were preferentially represented in NAT-associated niches (p<0.0001; **Fig. S12c**). The first, termed Mucosal macrophages, was enriched for phagocytosis and bacterial-response processes (FDR < 0.05; **Fig. S12d**). The second, termed Stromal macrophages, showed enrichment for macrophage activation and immunoregulatory signaling pathways (FDR < 0.05; **Fig. S12e**). In contrast, two macrophage states were enriched in CRC-associated niches (p<0.0001; **Fig. S12c**). First, *SPP1*+ macrophages were enriched for ECM remodeling pathways (FDR < 0.05; **Fig. S12f**) and preferentially localized to the CAF niche and lymphocyte-rich fibrotic stroma. Second, an *MMP12*+ state, which was not described in the original analysis of this dataset (16), was enriched for inflammatory signaling pathways (FDR < 0.05; **Fig. S12g**) and localized to inflammatory tumor-immune interface.

## Discussion

We introduced CIRCA, a self-supervised causal representation learning framework for jointly identifying cell- and microenvironment-level transcriptional programs and their interactions from spatial transcriptomics data. Grounded in theoretical identifiability results under assumptions about the underlying causal process, CIRCA enables the recovery of reproducible, interpretable latent factors without requiring predefined biological gene programs or interaction networks. Benchmarking across diverse spatial transcriptomics platforms demonstrated its flexibility, superior niche identification, and ability to recover higher-order tissue architecture from local cell-neighborhood relationships. Together, these results demonstrate how CIRCA provides a theoretically grounded framework for characterizing tissue architecture and modeling cell-microenvironment interactions.

Using spatial transcriptomics and Aβ pathology measurements from an AD mouse model, CIRCA revealed regional differences in microglial responses to Aβ plaques and identified interpretable transcriptional programs and their interactions with Aβ pathology. Consistent with previous analysis of this dataset (49), plaque-associated microglial responses were most concordant between the cortex and hippocampus. However, CIRCA enabled us to additionally identify distinct white matter and thalamic plaque niches that exhibited greater transcriptional heterogeneity. Despite these differences, microglia across regions exhibited a shared shift from homeostatic toward disease-associated states (51), with thalamic microglia showing weaker enrichment for classical DAM signatures and lower concordance with other regions across broader microglial state signatures. Interestingly, AD genetic risk signatures were also preferentially enriched in hippocampal and cortical plaque-associated microglia, consistent with previous regional associations identified using seismic (65). Together, these findings suggest that microglial responses to Aβ pathology and their association with AD genetic susceptibility depend on anatomical and pathological context, consistent with broader observations of regional heterogeneity in AD pathophysiology and progression (70, 71).

Beyond regional differences, CIRCA identified distinct transcriptional programs and their interactions with Aβ pathology at multiple spatial scales. Notably, cell-level Aβ burden was associated with two distinct DAM-like factors (51, 72, 73): one characterized by phagocytosis, lysosomal processing, and antigen presentation, and another by inflammatory signaling, suggesting that functional programs involved in plaque recognition and clearance can be distinguished from those associated with inflammatory responses. Cell-level Aβ burden was also linked to a disease-associated astrocyte-like program (67), while niche-level factors captured additional DAM-associated and immune-related transcriptional programs in the surrounding microenvironment. Interestingly, a niche factor linked to both cell- and neighborhood-level Aβ burden was also enriched for ERBB4 signaling, recently implicated in amyloid-associated neurodegeneration through aberrant activity in vulnerable excitatory neurons (74). Although this association requires further validation, it suggests a potential link between plaque-associated microenvironments and neuronal signaling pathways implicated in AD. Together, these findings highlight CIRCA’s ability to resolve multiple potentially interacting components of pathology-associated tissue dysregulation.

Additionally, applying CIRCA to a human tumor dataset (CRC) demonstrated its generalizability, revealing shared spatial organization of remodeled stromal microenvironments in the presence of substantial patient-specific heterogeneity among malignant populations. The conserved spatial relationships among these stromal niches suggest recurring patterns of stromal remodeling despite patient-specific differences in their spatial extent. This organization highlights the value of characterizing tumor microenvironments beyond proximity to malignant tissue, which was used to define the tumor microenvironment in the original analysis (16). These findings are consistent with previous studies identifying coordinated cellular neighborhoods and spatially organized tumor–stromal interfaces in CRC (75–77) and other cancers (78), some of which have been linked to prognosis or treatment response. Within these remodeled compartments, CIRCA further resolved distinct macrophage states. The localization of *SPP1*+ macrophages to CAF-associated and lymphocyte-rich fibrotic stromal niches agrees with prior work linking *SPP1*+ macrophages and fibroblasts to ECM remodeling and desmoplastic tumor microenvironments (79). In contrast, CIRCA identified an *MMP12*+ macrophage population concentrated at the inflammatory tumor–immune interface that was not resolved in the original analysis (16). Together with previous observations of spatially specialized macrophage populations in CRC (80), these findings suggest that distinct tumor-associated macrophage states may participate in different aspects of stromal remodeling and tumor-immune interactions.

Despite these strengths, CIRCA has several limitations. Although CIRCA is built on a framework with theoretical guarantees of latent factor and interaction identifiability, these guarantees rely on assumptions about the underlying data-generating process that may not be met in real tissue. Even when these assumptions hold, theoretical identifiability does not guarantee accurate recovery of latent factors and their interactions in finite samples. Consequently, CIRCA’s inferred interactions cannot definitively establish cause-effect relationships in real spatial transcriptomics data. Furthermore, systematic within-section technical variation may also be captured by CIRCA’s latent factors, although its factorized latent representations could facilitate the identification and exclusion of artifact-associated factors from downstream analyses. Nonetheless, empirical robustness to certain forms of model misspecification demonstrated in previous work (42), together with CIRCA’s strong benchmarking performance and factor reproducibility, supports the utility of its learned representations for biological analysis. Together, these findings suggest that CIRCA provides a robust framework for learning biologically informative latent representations from spatial transcriptomics data.

Our biological applications are subject to additional limitations. Partially due to the limited availability and high cost of cell-resolution spatial transcriptomics technologies, many available datasets, including those analyzed here, contain samples from relatively few individuals. This limits our ability to assess associations between learned factors or spatial niches and patient-level phenotypes, as well as the generalizability of the identified biological patterns. Additionally, our AD analysis was restricted to a mouse model because we sought to incorporate paired Aβ pathology measurements, which are not yet available in human cell-resolution datasets. Consequently, the identified plaque-associated transcriptional programs and their regional differences may not fully generalize to pathophysiology in human AD. Although such paired data is currently limited, advances in spatial multimodal profiling are making joint measurements of gene expression and tissue pathology increasingly accessible, providing opportunities to extend CIRCA to human disease.

In conclusion, we present CIRCA, a theoretically grounded framework for learning reproducible, interpretable representations of tissue architecture and cell-microenvironment interactions from spatial transcriptomics data. Across diverse tissues and disease contexts, CIRCA recovered shared patterns of tissue organization and distinct transcriptional programs associated with pathological remodeling, highlighting its potential for investigating the molecular processes underlying tissue organization and disease.

## Methods

### CIRCA

#### Input data

We define a spatial transcriptomic dataset as 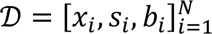, where *N* is the number of cells, *x_i_* is the raw gene expression count vector for cell *i*, *s_i_* are the spatial coordinates, and *b_i_* is the technical batch label [i.e. slide, field of view (FOV)]. CIRCA optionally incorporates additional omics layers or covariates *u_i_* so long as they can be mapped to the same cells in *D*, which are variance-stabilized if appropriate and standardized before model input.

#### Neighborhood graph

To represent local spatial relationships, we construct batch-specific k-nearest neighbor graphs G*_b_* = (V*_b_*, ℰ*_b_*) for all batches *b* ∈ *B* using Euclidean distances between cell coordinates. The global graph, G = (V, ℰ) is simply the union of all batch-specific graphs, ensuring that neighborhoods do not span batches. Each node *v_i_* ∈ V represents a cell and each edge *e_ij_* = (*v_i_*, *v_j_*) ∈ ℰ indicates that nodes *v_i_* and *v_j_* are spatial neighbors. A cell *i*’s local neighborhood N(*i*) is defined as the set of *k* cells [*v_j_*] where *e_ij_* = (*v_i_*, *v_j_*) ∈ ℰ.

#### Data augmentation

During training, CIRCA applies data augmentations to gene expression profiles and spatial neighborhoods to improve robustness to technical variation. To reduce dependence on library size, total counts are permuted among cells within each mini-batch, and expression profiles are rescaled to the corresponding permuted totals. To simulate dropout, expression profiles are randomly masked, downsampled, or both at a target rate of 20%. Finally, to improve robustness to segmentation errors and cell filtering, local neighborhoods are augmented by uniformly sampling *k* neighbors from the 1.2*k* nearest spatial neighbors, approximating a 20% dropout rate in local neighborhood membership.

#### Model overview

CIRCA adapts the recently developed causal representation learning framework, G-CaRL (42), to model latent cell- and niche-level transcriptional programs and their interactions in spatial transcriptomics data. For each cell *i*, we define a central cell expression group *c_i_* = *x_i_* and its local neighborhood expression group *n_i_* = [*x_j_*, ∀*j* ∈ N(*i*)]. We assume a data-generating process where observations are produced through group-specific mixing of interacting latent factors such that 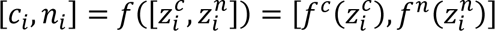, where 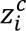 and 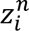 represent latent cell- and neighborhood-level transcriptional programs, respectively, and *f*, *f^c^*, and *f^n^* represent unknown mixing functions. This framework can be expanded to incorporate different feature groups, such as pathology measurements or additional omics modalities.

We model the interactions among these latent factors using the following pairwise factorization of their joint distribution, defined such that interactions between individual factors across groups resemble a pairwise Bayesian network. In this factorization, the joint distribution over the latent factors is given as

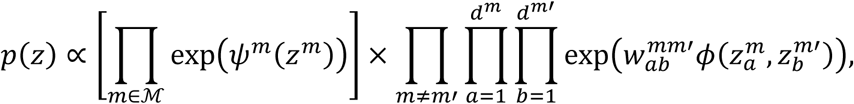

where *M* denotes the set feature groups, *d_m_* denotes the number of latent factors in group *m*, *ψ_m_* captures within-group dependencies, and *φ*(⋅,⋅) is a pairwise potential function representing a causal relationship between two latent factors. The coefficient 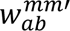 modulates the strength and sign of causal interactions between feature *a* in group *m* and feature *b* in group *m*’, forming the weighted causal adjacency matrices *W^mm^*^’^.

Under this framework, latent factors that interact across groups are identifiable up to permutation and component-wise invertible transformations, provided that the inter-group interaction structure is sufficiently nondegenerate and the pairwise potential function *φ*(⋅,⋅) satisfies conditions involving asymmetric, non-factorizable cross-derivatives. Identifiability of the interaction structure additionally requires directed interactions, connectivity through shared-parent or shared-child relationships, and stronger conditions on causal potential function, yielding recovery up to permutation, scaling, and matrix transposition. The formal assumptions and identifiability results are detailed in the **Supplementary Methods**.

When these assumptions are met, these latent representations and their interactions can be recovered using a self-supervised contrastive objective. CIRCA is trained to discriminate observed cell-neighborhood pairs from synthetic negatives generated by independently permuting feature groups. Specifically, positive and negative pairs are defined as 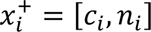 and 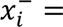 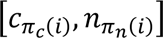, respectively, where *π_c_* and *π_n_* are independently sampled random permutations. CIRCA learns to distinguish these samples using a factorized nonlinear discriminator with the form

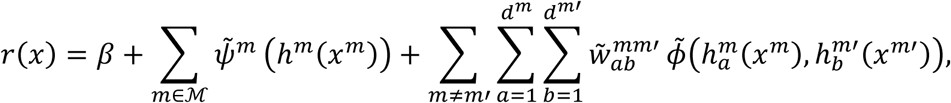

where ℎ*^m^* denotes a group-specific encoder, 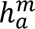 is the *a*-th factor of ℎ*^m^*, *β* is a bias term, and 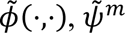, and 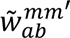 are learned parameters corresponding to the generative model defined above. The causal representation learning objective is

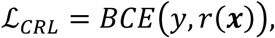

where BCE is the binary cross-entropy loss and *y* indicates whether a sample corresponds to an observed or permuted pair. Because groupwise permutations preserve the marginal distributions of each feature group while disrupting dependencies introduced by causal interactions between them, the discriminator, constrained to reflect the factorization of the assumed generative model, must recover the underlying latent representations and their interactions to optimally distinguish observed from permuted samples.

#### Model Implementation

CIRCA uses an encoder-only architecture to learn cell-state embeddings and identifiable cell- and niche-level latent factors through self-supervised contrastive objectives. The encoders are implemented as multilayer perceptrons (MLPs) consisting of fully connected layers with layer normalization, leaky ReLU activation functions, and dropout.

A shared MLP first embeds the augmented expression profiles of the central cell and its neighbors, then splits the embedding into cell (ℎ*_c_*) and neighborhood (ℎ*_n_*) components. While ℎ*_c_* is passed directly to the cell encoder, the neighborhood embeddings undergo an additional aggregation step. Mean- and max-pooled neighbor embeddings are each projected to lower-dimensional summaries and concatenated with low-dimensional projections of each neighbor, ordered by distance to the central cell, capturing both aggregate neighborhood characteristics and how proximity interacts with neighbor features. Finally, separate MLP encoders then produce the cell- and niche-level latent factors *Z^c^* and *Z^n^*, which are batch-normalized without learnable affine parameters to stabilize their scale and keep the magnitudes of learnable edges in *W* comparable. Additional feature groups, such as pathology, are encoded by their own MLPs.

#### Auxiliary loss term

While CIRCA’s identifiable cell embeddings capture spatially interacting cell information, they do not necessarily capture all cell state information and may be suboptimal for cell type identification. To this end, we add an additional MLP projection head to the model backbone that learns cell state embeddings useful for cell-typing alongside CIRCA’s spatially interacting latent factors using the self-supervised, contrastive SimCLR loss (81), given as

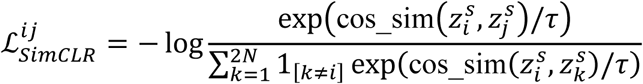

where 1_[*k*≠*i*]_is an indicator function, *z^s^* is a concatenated embedding matrix computed from two different augmented views of each cell, and *i*, *j*, *k* ∈ 2*N* are sample indices in *z^s^*. As this loss function does not directly constrain the interacting latent factors, the identifiability results described above hold so long as the shared backbone has sufficient capacity.

#### Regularization

Theoretically, the above self-supervised classification task is sufficient to recover interacting cell and niche latent factors under the above assumptions. In practice, we include regularization terms to encourage the models search space to conform with these assumptions. First, we include a symmetry penalty that encourages the causal function *φ*(⋅,⋅) to remain asymmetric, defined as a hinge loss

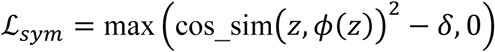

where cos_sim(*z*, *φ*(*z*)) indicates the cosine similarity between each latent factor *z* and its value after passing through the nonlinear causal function *φ*(⋅) and *δ* represents a collinearity threshold after which the penalty is applied.

We also apply a smooth approximation of an L_1_ regularization penalty on the causal adjacency matrix *W*, given as

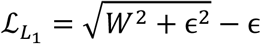

where *ε* is a small constant value. This helps to push the model towards capturing sparse, asymmetric interactions between groups.

Altogether, the combined loss function for CIRCA is given as

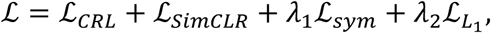

where *λ*_1_ and *λ*_2_ are regularization parameters determining the strength of the asymmetry and sparsity regularization terms. For detailed descriptions of hyperparameter settings, see the **Supplementary Methods**.

#### Model Interpretation

We use Integrated Gradients (45) with a zero-expression baseline to obtain signed gene-level attributions for each cell- and niche-level factor, aggregated across neighbors for niche factors and averaged across cells to give a gene-by-factor matrix. To reduce the dependence on mean expression, genes were binned into expression quantiles and attributions were divided by the root mean square of signed attributions pooled across factors within each bin, preserving attribution direction and relative contributions across factors. Inspired by SIGnature (82), we averaged these adjusted attributions over genes in curated pathway and biological process gene sets (83–87). Pathway attribution scores were tested and normalized against empirical null distributions generated from 10,000 gene-label permutations.

### Datasets

We used seven published cell-resolution spatial transcriptomics datasets (11, 12, 14, 46, 88, 89) to benchmark CIRCA and two additional datasets (16, 49) for biological applications. Dataset characteristics following preprocessing are summarized in **Table 1**, with detailed quality control and preprocessing procedures provided in the **Supplementary Methods**. Published cell type and spatial domain annotations were used as reference labels for benchmarking.

### Benchmarking

We benchmarked CIRCA against BANKSY (26), SEDR (23), GraphST (21), Novae (30), and NicheCompass (35) for spatial niche identification using seven spatial transcriptomics datasets with independent published anatomical domain annotations. Performance was evaluated across three dimensions: (1) biological accuracy (ARI, NMI, cLISI), (2) spatial continuity (FIDE, MLAMI), and (3) batch integration (kBET, iLISI). Metrics were min-max scaled within each dataset and averaged to create summarized scores for each dimension. The overall score is the mean of these scores. Each method was evaluated across five random seeds. Embeddings were clustered using Leiden clustering, with resolution optimized to match the number of annotated domains, except for Novae, which used its native assign_domains() function. Detailed metric definitions, model configurations, and benchmarking procedures are provided in the **Supplementary Methods** and **Supplementary Table S1**.

### Statistical analysis

Method comparisons between biological accuracy, spatial continuity, slide integration, and overall scores within datasets used Welch’s t-test with Holm correction (90). A mixed effects model with the formula Score ∼ Method + (1 | Dataset) + (1 | Method:Dataset) was used to compare performance in all benchmarking tasks combined. Differences in cell type proportions across niches were tested using Chi-square tests; heatmaps show the standardized residuals, which were tested post-hoc for each cell type-niche combination with Bonferroni correction (91). Unless stated otherwise, false discovery rates (FDR) were controlled using the Benjamini-Hochberg procedure (92).

#### Spatial transcriptomic analysis

The rapids-singlecell package (93) was used for constructing embedding nearest neighbor graphs, computing UMAPs, and Leiden clustering. PAGA graphs (47) were constructed with scanpy (94). Cell-level marker genes for cell types, cell states, and niches were identified with sc.tl.rank_genes_groups (method=’t-test_overestim_var’). Pseudobulk differential expression in AD mouse model microglia used pyDESeq2 (95, 96) on counts aggregated by tissue section and brain region, comparing plaque-associated niches with the corresponding non-plaque region (CTX, HIP, TH, and WM).

#### Pathway enrichment analysis

GSEA (97) was performed with GSEApy (98) on test statistics from the corresponding differential expression analysis using 10,000 permutations. Tumor-associated niches were tested for enrichment in Gene Ontology Biological Process (GO:BP) (83) terms with sizes within 15 to 500 genes, and plaque-associated microglia against published microglial subtype signatures (50).

#### Genetic association analysis

We analyzed GWAS summary statistics from 12 datasets covering early-onset AD (onset < 65 and < 70 years, respectively), AD, and AD and related dementias (53–62) (**Table S1**). Statistics were harmonized to human reference genome GRCh38 and converted to gene-level association statistics with MAGMA (99) using the SNP-wise mean model, a gene window of 35 kb upstream and 10 kb downstream, and the 1000 Genomes Phase 3 European data (100) as a reference to account for linkage disequilibrium. For each dataset, the top 1,000 genes by MAGMA z-score defined a GWAS-derived disease-associated gene set. We used scDRS (52) with default settings to compute cell-level disease relevance scores for each gene set and tested their association with cell types, niches, and anatomical region-stratified major cell types as described in (52). Because the CosMx gene panel is small, the number of mean and variance bins was automatically reduced from 20 to 9. For full details of GWAS summary statistic harmonization and MAGMA association analysis, see the **Supplementary Methods**.

## Data availability

All spatial transcriptomics datasets used in this study are publicly available. The BaristaSeq, MERFISH, osmFISH, and STARmap datasets are available at SDMBench (http://sdmbench.drai.cn/). The Allen Brain Cell Atlas datasets are available at the Allen Institute (https://alleninstitute.github.io/abc_atlas_access/descriptions/Zhuang_dataset.html). The mouse embryo dataset is available at STOmicsDB (https://db.cngb.org/stomics/mosta/). The AD mouse model CosMx dataset is available at Gene Expression Omnibus (GEO, accession number GSE263791). The Visium HD colorectal cancer dataset is available through 10x Genomics, Inc., at https://www.10xgenomics.com/platforms/visium/product-family/dataset-human-crc.

## Code availability

The source code for CIRCA, including model implementation, training, and analysis scripts, is publicly available at https://github.com/bsml320/CIRCA.

## Materials availability

This study did not generate new materials.

## Acknowledgements

This study was partially supported by grants from the National Institutes of Health (NIH, U01AG079847 and R01LM012806). We acknowledged technical support from the UTHealth Cancer Genomics Core funded by the Cancer Prevention and Research Institute of Texas (CPRIT RP240610). TCL is supported by a postdoctoral training fellowship from the Gulf Coast Consortia, on the NLM Training Program in Biomedical Informatics & Data Science (T15LM007093). NE received Ranajit Chakraborty, PhD Fellowship in Human Genetics. The funders had no role in study design, data collection, data analysis, interpretation, decision to publish, or preparation of the manuscript.

## Author contributions

T.C.L. conceived the project, developed the CIRCA method, and performed most analyses. C.T contributed to the biological interpretation of the AD and CRC results. N.E. performed GWAS analyses and contributed to their interpretation. Z.Z. supervised the project. All authors contributed to writing and reviewing the manuscript.

## Declaration of generative AI and AI-assisted technologies in the writing process

During the preparation of this work, the authors used ChatGPT and Claude to improve manuscript writing. After using this tool, the authors reviewed and edited the content as needed and take full responsibility for the content of the published article.

## Supplementary information

Document S1. Supplementary Materials: Table S1, Figures S1-12.

Document S2. Supplementary Methods.

Document S3. Supplementary Table S2.

