## Supplementary Methods for "CIRCA learns identifiable cell- and niche-level programs and their interactions with tissue pathology from spatial transcriptomics"

### CIRCA model overview

**Input data**. We define a spatial transcriptomic dataset as $\mathcal{D} = [x_i, s_i, b_i]$ for $i \in N$, where $N$ is the number of cells, $x_i$ is the transcriptomic feature vector of cell $i$, $s_i$ are the spatial coordinates of cell $i$, and $b_i$ is the batch label (i.e. slide, field of view (FOV)) for cell $i$, where applicable. Optionally, CIRCA can operate on additional omics layers or covariates $y_i$ that are mapped to the same cells in $D$. For unimodal data, $x_i$ consists of raw gene expression counts. In multimodal data, additional omics layers or covariates $u_i$ are variance stabilized if required, standardized, and formatted so they can be retrieved for each cell $i \in N$. We denote the corresponding matrices across all cells with capital letters, for example, $X = [x_0, x_1, \dots, x_N]$

**Neighborhood graph.** To represent spatial associations between a cell and its local neighborhood, we construct batch-specific k-nearest neighbor graphs $\mathcal{G}_b = (\mathcal{V}_b, \mathcal{E}_b)$ for all unique batches $b \in B$ based on Euclidean distances between spatial coordinates. The global spatial k-nearest neighbor graph, $\mathcal{G} = (\mathcal{V}, \mathcal{E})$ is simply the union of all batch-specific graphs. Each node $v_i \in \mathcal{V}$ represents a cell and each edge $e_{ij} = (v_i, v_j) \in \mathcal{E}$ indicates that nodes $v_i$ and $v_j$ are spatial neighbors. A cell $i$'s local neighborhood is defined as the set of $k$ cells $[v_j]$ where $e_{ij} = (v_i, v_j) \in \mathcal{E}$.

**Data augmentation.** Data augmentation is a technique to introduce artificial variation to data during training to improve the robustness and generalization of the model. In this work, we design biologically relevant augmentations to cell expression profiles and spatial neighborhood graphs for spatial transcriptomics data.

First, to encourage the independence of learned representations from cell library size, we shuffle the total counts of each cell in a batch and scale their counts to sum to the new permuted total. Second, to improve robustness to dropout, we artificially mask, down sample, or mask and down sample expression profiles at a rate of 20%.

To improve model robustness to low-quality segmentation and cell filtering, we also augment membership in each cell's local neighborhood as defined by the batch-specific k-nearest neighbors graphs described above. During training, we construct a nearest neighbor graph with the $1.2k$-nearest neighbors, then sample $k$ neighbors at a time uniformly at random, simulating a 20% dropout rate of cells in each local neighborhood.

**Model overview.** CIRCA adapts the recently developed causal representation learning framework, G-CaRL (1), to model cell-niche interactions in spatial transcriptomics data. The model leverages prior knowledge about the grouping of variables (*i.e.*, cell expression and local neighborhood expression) and pairwise constraints on the causal interactions between each group to identifiably learn both the unobserved latent factors and the causal graph of interactions between groups. Specifically, for a cell gene expression profile $c_i = [x_i]$ and a local neighborhood expression profile $n_i = [x_j \ \forall \ e_{ij} \in \mathcal{E}]$, we assume a data generating process such that $[c_i, n_i] = f([z_i^c, z_i^n]) = [f^c(z_i^c), f^n(z_i^n)]$, where $f$, $f^c$, and $f^n$ represent unknown mixing functions that transform the latent transcriptional programs $z_i^c, z_i^n$ into the observed gene expression vectors $c_i, n_i$, respectively.

Additionally, we assume that the interactions between latent transcriptional programs in a cell and its local neighborhood interact in a manner similar to pairwise Bayesian Networks and can be factorized as

$$p(z) \propto \left[\prod_{m\in M} exp(\psi^m(z^m))\right] \times \prod_{m\neq m'} \prod_{(a,b)\in|z^m|\times|z^{m'}|} exp\left(w_{ab}^{mm'} \phi(z_a^m, z_b^{m'})\right),$$

where $M$ denotes the set of all groups in the model (the cell $c$, neighborhood $n$, and any additional covariates or omics layers $u$), $\phi(\cdot,\cdot)$ denotes a potential function representing a causal relationship between two latent variables, $\psi^m$ denotes a group-wise potential function representing any causal relationships within a group $m$, and the coefficient $w_{ab}^{mm'}$ modulates the strength of the causal interaction between feature $a$ in group $m$ and feature $b$ in group $m'$. The matrix $W^{mm'} = \left(w_{ab}^{mm'}\right)_{a\in|z^m|,b\in|z^{m'}|}$ can be interpreted as a weighted adjacency matrix for causal interactions between group $m$ and $m'$.

Under the grouped observational model and pairwise latent-variable factorization described above, (1) established the following conditions for identifying latent variables and their inter-group causal relationships.

**Latent factor identifiability (Theorem 1).** The latent variables within each group are identifiable up to permutation and variable-wise invertible transformations under two assumptions:

1. Nondegeneracy of the causal graph (A1): For any group $m$ in $M$, each latent variable has at least one neighbor in another group, and the collection of inter-group adjacency matrices has full row-rank after removing all-zero rows.
2. Causal function (A2): The pairwise causal function $\phi(\cdot,\cdot)$ must be asymmetric and have non-factorizable cross-derivatives.

The first assumption is relatively straightforward and simply requires that the latent variables in each group interact with at least one other group, and that the causal strengths of those interactions differ for both causal directions. The second assumption requires that the causal function be asymmetric with regards to the order of the latent factors and sufficiently complex that its cross-derivative cannot be factored, enabling the model to distinguish one causal direction from another.

**Inter-group causal graph identifiability (Proposition 2).** Additional assumptions are required for the causal graph of interactions between groups to be identified up to permutation, linear scaling, and matrix transpose:

3. Causal graph structure (C'1): All inter-group causal relationships are directed, and for every group $m$, all variables in that group have either a shared cause or shared effect with another variable in group $m$. Furthermore, all variables within each group must be connected.
4. Causal function (C'3): In addition to the constraint on the pairwise causal function $\phi(\cdot,\cdot)$ described above, we exclude causal functions whose cross-derivative $\phi^{12}(\cdot,\cdot)$ can be decomposed as:

$$\phi^{12}(x,y) = \alpha(x)\beta(y)\exp(\gamma(x,y) - \gamma(y,x)))$$

This assumption is a stricter requirement on the structure of the causal graph than the first assumption above, requiring that each latent variable in a group $m$ has another factor in $m$ with a shared cause or shared effect in another group. However, we note that even when this condition fails, the latent factors themselves can still be recovered so long as assumptions 1 and 2 are met.

When these conditions are met, the latent factors $Z$ and the causal adjacency matrix $W$ can be jointly recovered using a self-supervised contrastive learning framework. CIRCA is trained to discriminate real cell-neighborhood pairs from permuted ones using a nonlinear logistic regression system that factorizes according to the pairwise Bayesian network described above. Formally, we classify $\boldsymbol{x_N} = [c_{i\in N}, n_{i\in N}]$ vs $\boldsymbol{x_{N^*}} = [c_{i\in N^*}, n_{i\in N^*}]$ by minimizing the binary cross-entropy loss given by a regression function with the form

$$r(\boldsymbol{x}) = c + \sum_{m\in M} \psi^m\big(h^m(x^m)\big) + \sum_{m\neq m'} \sum_{(a,b)\in|z^m|\times\left|z^{m'}\right|} w_{ab}^{mm'} \, \phi\left(h_a^m(x^m), h_b^{m'}\left(x^{m'}\right)\right)$$

where $h^m$ denotes a group-wise encoder feature extractor, $h_a^m$ is the $a$-th element of $h^m$, $c$ is a bias term, and $\phi(\cdot,\cdot)$, $\psi^m$, and $w_{ab}^{mm'}$ are defined as above. This gives our causal representation learning loss:

$$\mathcal{L}_{CRL} = BCE\big(y, r(\boldsymbol{x})\big)$$

where BCE is the binary cross-entropy loss and $y$ is the label indicating whether a sample in $x$ corresponds to a real data point or a permuted one.

It is not immediately obvious why solving this binary classification task enables the recovery of the underlying causal structure. However, the group-wise permutations applied to the input $x$ breaks the causal relationships between groups, requiring the above logistic regression problem to learn the inter-group causal relationships in the latent space to solve the classification task. Due to the set-up of the regression model matching the factorization of the data-generating process above, the optimal model is only achieved when the ground truth is recovered up to the transformations described above.

**Model Implementation.** CIRCA uses an encoder-only architecture to learn cell-state embeddings and identifiable cell- and niche-level latent factors through self-supervised contrastive objectives. The encoders are implemented as multilayer perceptrons (MLPs) consisting of fully connected layers with layer normalization, leaky ReLU activation functions, and dropout. Each hidden layer $h^{(l)}$ is defined as

$$h^{(l)} = \text{Dropout}\left(\text{LeakyReLU}\left(\text{LayerNorm}\left(A_l h^{(l-1)} + b_l\right)\right)\right),$$

where $h^{(l-1)}$ is the previous hidden layer and $A_l$ and $b_l$ correspond to the weights and biases of the $l^{\text{th}}$ linear layer, respectively.

CIRCA first embeds the augmented expression profiles of the central cell and its spatial neighbors into a lower-dimensional embedding space using a shared MLP. The resulting embeddings are separated into cell and neighborhood components, $h_c$ and $h_n$, which are subsequently processed by separate MLP encoders to obtain the cell- and niche-level latent factors, $Z^c$ and $Z^n$. While $h_c$ is passed directly to the cell encoder, the neighborhood embeddings undergo an additional projection and aggregation step. Specifically, mean- and max-pooled neighborhood embeddings are independently projected into lower-dimensional summaries, which are then concatenated with shared low-dimensional projections of each neighbor, ordered by spatial proximity to the central cell. This approach enables CIRCA to capture both aggregate neighborhood characteristics and information about the interaction between spatial proximity and neighbor-specific features. Finally, batch normalization without learnable affine parameters is applied to the latent factors before computing the factorized discrimination loss, stabilizing their scale and ensuring the magnitude of learnable edges in the causal adjacency matrix $W$ are comparable. Any additional feature groups, such as pathology information, are encoded with additional MLPs that directly compute the corresponding latent factors.

**Auxiliary loss term.** While CIRCA's identifiable cell embeddings capture spatially interacting cell information, they do not necessarily capture all cell state information and may be suboptimal for cell type identification. To this end, we add an additional MLP projection head to the model backbone that learns cell state embeddings alongside CIRCA's spatially interacting latent factors using the self-supervised, contrastive SimCLR loss (2), given as

$$\mathcal{L}_{SimCLR}^{ij} = -\,log\frac{exp\left(\mathrm{cos\_sim}\left(z_i^s, z_j^s\right)/\tau\right)}{\sum_{k=1}^{2N} \mathbb{1}_{[k\neq i]}\, exp\left(\mathrm{cos\_sim}\left(z_i^s, z_k^s\right)/\tau\right)}$$

where $\mathbb{1}_{[k\neq i]}$ is an indicator function, $z^s$ is a concatenated embedding matrix computed from two different augmented views of each cell, and $i, j, k \in 2N$ are sample indices in $z^s$. This loss term ensures that CIRCA does not ignore information that is not strongly spatially varying early in training and provides an embedding space not dependent on spatial location that is useful for cell type identification. Additionally, as this loss function does not directly influence the interacting latent factors, its inclusion does not harm the identifiability results described above so long as the shared backbone has sufficient capacity.

**Regularization.** Theoretically, the above self-supervised classification task is sufficient to recover interacting cell and niche latent factors under the above assumptions. In practice, we include regularization terms to encourage the models search space to conform with these assumptions. First, we include a symmetry penalty that encourages the causal function $\phi(\cdot,\cdot)$ to remain asymmetric, defined as a hinge loss

$$\mathcal{L}_{sym} = \max\left(\mathrm{cos\_sim}\left(z, \phi(z)\right)^2 - \delta, 0\right)$$

where $\cos\_\mathrm{sim}(z, \phi(z))$ indicates the cosine similarity between each latent factor $z$ and its value after passing through the nonlinear causal function $\phi(\cdot)$ and $\delta$ represents a collinearity threshold after which the penalty is applied.

We also apply a smooth approximation of an $L_1$ regularization penalty on the causal adjacency matrix $W$, given as

$$\mathcal{L}_{L_1} = \sqrt{W^2 + \epsilon^2} - \epsilon$$

where $\epsilon$ is a small constant value. This helps to push the model towards capturing sparse, asymmetric interactions between groups.

Altogether, the combined loss function for CIRCA is given as

$$\mathcal{L} = \mathcal{L}_{CRL} + \mathcal{L}_{SimCLR} + \lambda_1 \mathcal{L}_{sym} + \lambda_2 \mathcal{L}_{L_1},$$

where $\lambda_1$ and $\lambda_2$ are regularization parameters determining the strength of the asymmetry and sparsity regularization terms.

**Model Interpretation.** While CIRCA recovers an identifiable latent space through the causal representation learning procedure described above, the model is an encoder-only setup and requires post-hoc interpretation techniques to link gene expression features to the recovered latent factors. To do this, we use the Integrated Gradients (IG) method (3). For a given function $F$, input sample $x_i$, and baseline vector $x'$, the IG method calculates the contribution of each feature to an output value by computing a path integral over the gradients of the model along the line from the baseline to the input sample:

$$IG(x_i, x', F) = (x_i - x') \times \int_{\alpha=0}^{1} \frac{\partial F\left(x' + \alpha \times (x_i - x')\right)}{\partial x_i} d\alpha$$

The IG method has two key properties that make it useful for interpreting the model learned by CIRCA. First, the model can efficiently map the contributions of all genes simultaneously to any real output of the model, including each of the latent factors and the logits for discriminating between real and permuted cell-neighborhood pairs. Second, the completeness axiom of the IG method states that the sum of all feature attributions adds up to the difference of the value of the model output at sample $x_i$ and the baseline $x'$. This allows us to sum gene-level attributions from all cells in the local neighborhood together and get a valid set of attributions for gene expression in the local neighborhood.

We use this IG approach to map gene-level contributions from cell expression to CIRCA's cell latent factors and from local-neighborhood expression to its niche latent factors. A zero-expression baseline was used, such that the sign of each attribution reflects whether expression of a given gene increases or decreases the corresponding latent factor. For each dataset, we averaged the signed IG attributions across cells to obtain a gene-by-latent-factor attribution matrix. Because IG magnitude is strongly influenced by baseline gene expression, highly expressed genes tend to receive larger absolute attributions. To account for this dependence, we bin genes into expression quantiles and divide attributions by the root mean square of signed attributions pooled across factors within each bin. This reduces the expression-dependent differences in attribution magnitude while preserving attribution direction.

Inspired by SIGnature (4), we also use our signed IG results to identify known pathways and biological processes enriched in each latent factor. Based on gene sets derived from prior knowledge databases (5-9), we average our scaled IG attribution results from above over all genes in each gene set to score their activity in each latent factor. To normalize these pathway activity estimates and compute statistical significance, we perform permutation testing by permuting the gene labels in the gene-by-factor attribution matrices 10,000 times.

**Latent factor alignment.** To assess the reproducibility of the recovered latent factors, we train multiple models with different random initializations of encoder weights and train-validation splits. Once each model has converged, we compute the Spearman cross-correlation matrix for all pairs of latent factors within each group on a set of cell-neighborhood pairs held out from the training of all models. As the latent factors are identified up to permutation, sign changes, and variable-wise transformations, we use scipy.optimize.linear_sum_assignment() to find the permutation that maximizes the sum of the absolute Spearman correlation along the diagonal of the cross-correlation matrix to identify which latent factors in one model correspond to factors in another. In cases where we want to set a lower bound on the absolute Spearman correlation for a latent factor to be considered replicated, we zero any entries of the cross-correlation matrix whose absolute value is below that threshold ahead of solving the linear sum assignment problem. Assigned pairs that are below this threshold are not considered to be replicated.

**Graph construction.** In addition to identifiable latent factors, CIRCA identifies the causal model of the latent causal interactions when the underlying assumptions are met. To convert the dense causal adjacency matrix $W$ estimated during training into a graphical model, we perform the following procedure: first, as the interactions are assumed to be directed, we compare weights representing opposing directed interactions ($w_{ij}$ vs. $w_{ji}$) in the adjacency matrix $W$ based on their absolute value and set the smaller values to zero, obtaining a fully connected directed graph. Next, we would like to prune edges in this directed graph to construct a sparse interaction model. To do this, we construct graphs across a range of sparsity values using hard thresholding and modify the trained CIRCA model to replace the dense matrix $W$ with the thresholded matrices $\widehat{W}$ representing directed graphs of increasing sparsity. We evaluate the AUROC of each of these models in the training task of discriminating real versus permuted cell-neighborhood pairs. As expected, we observe this performance generally decay as the sparsity increases along the range of thresholding values. To select a sparse graphical model that does not excessively prune true interactions, we identify the elbow point in this sparsity versus AUROC curve as the point with the greatest perpendicular distance from a linear interpolation between the lowest and highest threshold. Finally, we use that threshold to construct the final graphical model learned by CIRCA.

**Hyperparameter selection.** Training CIRCA models involves the selection of hyperparameters such as the size of the latent dimension and spatial neighborhoods, as well as regularization parameters such as the $L_1$ regularization strength. As the models learned by CIRCA are most sensitive to latent dimension and neighborhood size, we performed a grid search over latent dimensions of [25, 50, 75, 100] and neighborhood sizes of [5, 10, 15] for the models learned in the AD mouse model dataset and the Visium HD CRC dataset. To assess model quality, we aligned the latent factors of models trained on three different random seeds using the procedure described above and selected the models with the highest mean correlation coefficient (MCC) over all aligned factors and for each pairwise combination of models. Key hyperparameters for models included in the manuscript, including Latent dimension $d$, spatial neighborhood size $k$, symmetry regularization strength $\lambda_1$, and $L_1$ regularization strength $\lambda_2$, are as follows:

- **BaristaSeq:** $d = 10, k = 10, \lambda_1 = 0.02, \lambda_2 = 0.01$
- **MERFISH:** $d = 10, k = 10, \lambda_1 = 0.02, \lambda_2 = 0.01$
- **osmFISH:** $d = 10, k = 10, \lambda_1 = 0.02, \lambda_2 = 0.01$
- **STARmap:** $d = 10, k = 10, \lambda_1 = 0.02, \lambda_2 = 0.01$
- **Allen Brain Cell Atlas:** $d = 50, k = 10, \lambda_1 = 0.004, \lambda_2 = 0.002$

- **Mouse Embryo:** $d = 100$, $k = 10$, $\lambda_1 = 0.002$, $\lambda_2 = 0.001$
- **AD mouse model:** $d = 50$, $k = 10$, $\lambda_1 = 0.002$, $\lambda_2 = 0.001$
- **Colorectal cancer:** $d = 50$, $k = 10$, $\lambda_1 = 0.002$, $\lambda_2 = 0.001$

### Datasets

We used seven published spatial transcriptomics datasets to benchmark CIRCA and two additional datasets for biological applications. Dataset characteristics are depicted in **Table 1**. Published cell type and spatial domain annotations were used as ground truth for benchmarking.

**BaristaSeq.** Downloaded from SDMBench (10), this mouse visual cortex dataset (11) contained 5,527 cells across three sections and 79 genes. Excluding cells with fewer than 10 counts and genes expressed in fewer than 20 cells yielded 4,754 cells and 77 genes.

**MERFISH.** Downloaded from SDMBench (10), this mouse hypothalamus dataset (12) contained 28,317 cells across five consecutive sections and 155 genes. Removing cells in the bottom $5^{th}$ percentile of total counts yielded 26,901 cells.

**osmFISH.** Downloaded from SDMBench (10), this mouse somatosensory cortex dataset (13) contained 4,839 cells and 33 genes from a single tissue section.

**STARmap.** Downloaded from SDMBench (10), this mouse medial prefrontal cortex dataset (14) contained 3,190 cells across three sections and 166 genes.

**Allen Brain Cell Atlas (MERFISH).** We constructed two datasets from sequential sections of the Allen Brain Cell Atlas (15): eight coronal sections from mouse 1 (slices 1.074–1.082) and three sagittal sections from mouse 3 (slices 3.009–3.011). Following the original publication, we retained only cells with high-confidence reference-based annotations. We additionally excluded cells lacking spatial coordinates or with total counts below the $1^{st}$ percentile (21 counts for coronal; 34 for sagittal). The resulting coronal and sagittal datasets contained 239,857 and 239,795 cells, respectively, measured across 1,122 genes.

**Mouse Embryo (Stereo-seq)**. This dataset (16) contained 281,377 cells and 28,103 genes. We reversed the original log normalization to recover counts and retained the 6,000 most highly variable genes identified using scanpy.pp.highly_variable_genes() (17). Published cell type annotations were used for benchmarking. Spatial domain annotations were transferred from the corresponding bin50 Stereo-seq data to the cellbin data based on spatial proximity after coordinate alignment with STAlign (18).

**AD Mouse Model (CosMx).** This CosMx 1K dataset (19) contained 43,117 cells measured across 950 genes from four hippocampal sections of a humanized-APP AD mouse model, with paired cell-level amyloid-β (Aβ) immunofluorescence intensities. To mitigate previously reported nonspecific probe binding to Aβ plaques, we excluded genes whose expression correlated with at least one negative probe (Pearson's $r > 0.2$). We also calculated cell-level signal-to-noise ratios (SNRs) using the negative control probes and excluded cells with SNR < 3, fewer than 200 total counts, or Scrublet doublet scores > 0.2 (sc.pp.scrublet()) (20) to remove low-quality cells and potential segmentation errors. The final dataset contained 36,455 cells and 900 genes. Cell type and niche annotations were manually assigned following Leiden clustering of CIRCA's cell-state and niche embeddings, respectively.

**Colorectal Cancer (Visium HD).** This dataset consisted of three colorectal cancer (CRC) and two normal adjacent tissue (NAT) sections downloaded from 10x Genomics, Inc. (21). Cell-level counts were obtained using bin2cell (22) to segment nuclei and assign 2- µm bins to cells, yielding 1,136,534 cells measured across 17,996 genes. We excluded cells with fewer than 100 counts or greater than 30% mitochondrial counts and retained the union of the top 5,000 highly variable genes identified separately in each section using scanpy.pp.highly_variable_genes(). The final dataset contained 937,559 cells and 9,472 genes. Cell type and niche annotations were manually assigned following Leiden clustering of CIRCA's cell-state and niche embeddings, respectively.

**GWAS summary statistics.** Genome-wide association study (GWAS) summary statistics for Alzheimer's disease (AD) and early-onset Alzheimer's disease (EOAD) were sourced from previously published studies and publicly available genetic association repositories. A total of 12 GWAS summary-statistic datasets were included to evaluate gene-level associations across different AD phenotypes, age-at-onset definitions, and study populations.

EOAD summary statistics corresponding to age-at-onset thresholds of $< 65$ years and $< 70$ years obtained from (23). Additional early-onset AD association statistics were obtained from FinnGen, with the underlying biobank resource described by (24).

For AD and Alzheimer's disease and related dementias (ADRD), GWAS summary statistics were derived from the 2026 consensus meta-analysis (25), (26), (27), (28), (29), and (30). Additional datasets included FinnGen-derived AD association statistics (24), Pan-UK Biobank-derived AD association statistics (31), and the Alzheimer's Disease Genetics Consortium (ADGC) multi-ancestry GWAS available through the National Institute on Aging Genetics of Alzheimer's Disease Data Storage Site (NIAGADS; accession NG00177) (32).

**GWAS summary-statistics harmonization.** GWAS summary statistics were prepared for subsequent gene-based association analysis by utilizing SNP-level association p-values, genomic coordinates, variant identifiers, and sample-size information. The reference genome assembly was standardized to GRCh38 (hg38) to ensure consistency between variant positions and gene annotations.

Input files were formatted for MAGMA analysis using the available SNP identifiers, association p-values, and corresponding study-level or variant-specific sample sizes. For case-control GWAS, effective sample sizes were calculated based on the number of cases and controls, where applicable. Genomic coordinates and reference annotations were required to correspond to the same genome assembly.

### Benchmarking

To evaluate the performance of CIRCA for spatial niche identification, we compared CIRCA against BANKSY (33), SEDR (34), GraphST (35), Novae (36), and NicheCompass (37) in datasets with independently annotated spatial domains to serve as the ground truth. We evaluate performance across three key factors: biological accuracy, spatial continuity, and batch integration. All methods were run after initialization with 5 different random seeds to capture variability in model performance. All methods other than Novae were clustered with Leiden clustering, while Novae was clustered via their assign_domains() function.

**Biological accuracy metrics.** To assess the biological accuracy of the embedding spaces learned by each method, we first clustered the embedding space into the number of classes in the ground truth domain annotation using Leiden clustering with a binary search over resolution values in the range of 0 to 2. After each embedding space is clustered, we computed the Adjusted Rand Index (ARI) and Normalized Mutual Information (NMI) between the Leiden clustering labels and the ground truth domain annotations. We also computed the cell-type Local Inverse Simpson's Index (CLISI) over the 15-nearest neighbors in the embedding space using the spatial domain annotations as the cell type label. All three values are maximized when the embedding space captures the information in the ground truth labels. Metrics were min-max scaled within each dataset and averaged to create a summarized score for biological accuracy.

**Spatial continuity metrics.** To assess the spatial continuity of niches identified by the unsupervised clustering described above, we used the F1-score of inter-domain edges (FIDE) (36) and Maximum Leiden Adjusted Mutual Information (MLAMI) (37) scores. The FIDE score computes the F1-score for matching spatial domains over each cell's 15-nearest spatial neighbors. The MLAMI score computes Leiden clustering's of both the embedding space and the spatial coordinates over a range of resolutions and reports the configuration that gives the maximum AMI. Both values are maximized when spatial domains are contiguous. Metrics were min-max scaled within each dataset and averaged to create a summarized score for spatial continuity.

**Batch integration metrics.** To assess the ability of each approach to integrate data across different tissue sections, we computed the k-nearest-neighbor batch-effect test (kBET) and integration Local Inverse Simpson's Index (iLISI) scores. The kBET score examines the 15 nearest neighbors in each embedding space and tests whether the observed batch frequencies differ significantly from the expected global frequences. The iLISI score is also computed over the 15-nearest neighbors in the embedding space using the batch labels. Both values are maximized when batches are well-mixed. Metrics were min-max scaled within each dataset and averaged to create a summarized score for batch integration.

**Overall performance.** To create a balanced score capturing the overall performance of each method for the task of spatial domain identification, we took the mean of the summarized scores described above. In datasets that did not include multiple tissue sections (osmFISH and Mouse Embryo), only the biological accuracy and spatial continuity scores were averaged.

### Statistical analysis

Statistical significance between biological accuracy, spatial continuity, slide integration, and overall scores was computed using Welch's t-test, with the Holm method used for multiple hypothesis correction (38). A mixed effects model with the formula Score ~ Method + (1 | Dataset) + (1 | Method:Dataset) was used to compare performance in all benchmarking tasks combined. Tests for significance of differences of cell type proportions across niches or subsets of niches were computed using the Chi-square test, and the enrichment values displayed in heatmaps are the standardized residuals of the Chi-square tests. These values were used to assess significance of enrichment for a given cell-niche combination post-hoc, with the Bonferroni method used for multiple hypothesis correction (39).

**Spatial transcriptomic analysis.** The rapids-singlecell package (21) was used for constructing nearest neighbor graphs, computing UMAPs, and performing Leiden clustering in all datasets. PAGA graphs (40) were constructed with the sc.tl.paga() function in scanpy (17). Differential expression results computed at the cell-level, such as those used to identify cell type, cell state, and niche marker genes, were computed with sc.tl.rank_genes_groups() with method='t-test_overestim_var'. Differential expression results computed at the pseudobulk level were performed with pyDESeq2 (41, 42) after aggregating cell counts stratified by tissue section and the condition being tested.

**Pathway analysis.** Traditional GSEA analyses (43) were performed using the test statistic for the corresponding differential expression analysis using 10,000 permutations in GSEApy (44). To characterize biological processes enriched in tumor-associated niches, we tested for enrichment in Gene Ontology: Biological Process (GO:BP) (5) gene sets with at least 15 genes and fewer than

500 genes. To characterize transcriptional heterogeneity of microglia responses to amyloid-β plaques, we directly used a curated collection of gene signatures from previously published microglia subtypes (45).

**MAGMA gene-based association analysis.** Gene-based association analyses were conducted using Multi-marker Analysis of GenoMic Annotation (46) to aggregate SNP-level GWAS association evidence into gene-level association statistics. Genes were annotated according to gene coordinates aligned with the GRCh38 reference genome. SNPs were assigned to genes based on their genomic positions, with gene boundaries extended by 35 kilobases (kb) upstream and 10 kb downstream of each gene, accounting for gene orientation. Gene-based association statistics were calculated using the MAGMA SNP-wise mean model, which combines association signals from SNPs mapped to each gene while accounting for linkage disequilibrium (LD) among variants. LD was estimated using the European-ancestry (EUR) reference samples from the 1000 Genomes Project Phase 3 (47).

For each GWAS dataset, SNP-level association p-values and sample-size information were provided as input to MAGMA. The software evaluated the joint association evidence of variants mapped to each gene, accounting for their correlation structure, and generated an aggregate gene-level association p-value. Analyses were performed separately for all 12 GWAS datasets, generating a standardized set of gene-level association statistics for each AD or EOAD phenotype. The top 1000 genes based on z-score were used to define disease-associated gene sets for scDRS analyses (48).

**Genetic association analysis.** We used scDRS (48) with default settings to compute cell-level disease relevance scores for each gene set and tested their association with cell types, niches, and anatomical region-stratified major cell types as described in (48). Because the CosMx gene panel is small, the number of mean and variance bins was automatically reduced from 20 to 9. The null distribution was sampled with n_ctrl equal to 1,000.
