## Supplementary Materials for "CIRCA learns identifiable cell- and niche-level programs and their interactions with tissue pathology from spatial transcriptomics"

**Supplementary Table S1:** Characteristics of genome-wide association study summary statistics used in this work. Abbreviations: AD, Alzheimer's disease; ADRD, Alzheimer's disease and related dementias; EOAD, early-onset Alzheimer's disease.

| Dataset | Disease | AD proxy | # Case | # Control | # Samples | Reference |
|---|---|---|---|---|---|---|
| **AD_Jansen** | ADRD | Yes | 71,880 | 383,378 | 455,258 | (Jansen et al., 2019) |
| **AD_Kunkle** | ADRD | No | 35,274 | 59,163 | 94,437 | (Kunkle et al., 2019) |
| **AD_Schwartz** | ADRD | Yes | 53,689 | 355,900 | 409,589 | (Schwartzentruber et al., 2021) |
| **AD_Wightman** | ADRD | No | 90,338 | 1,036,225 | 1,126,563 | (Wightman et al., 2021) |
| **AD_Bellenguez** | AD | Yes | 111,326 | 677,663 | 788,989 | (Bellenguez et al., 2022) |
| **AD_Finngen** | AD | No | 16,652 | 483,534 | 500,186 | (Kurki et al., 2023) |
| **AD_2026** | ADRD | Yes | 128,681 | 849,833 | 978,514 | (Bellenguez et al., 2026) |
| **AD_NG00177** | AD | No | 17,839 | 19,543 | 37,382 | (Rajabli et al., 2025) |
| **AD_UKB** | AD | No | 839 | 410,833 | 411,672 | (Karczewski et al., 2025) |
| **EOAD_Age65** | EOAD | No | 3,182 | 13,386 | 16,568 | (Bradley et al., 2025) |
| **EOAD_Age70** | EOAD | No | 6,282 | 13,386 | 19,668 | (Bradley *et al.*, 2025) |
| **EOAD_Finngen** | EOAD | No | 1,965 | 191,784 | 193,749 | (Kurki *et al.*, 2023) |

**Supplementary Table S2:** Pathway-level attribution results for niche-level latent factors.

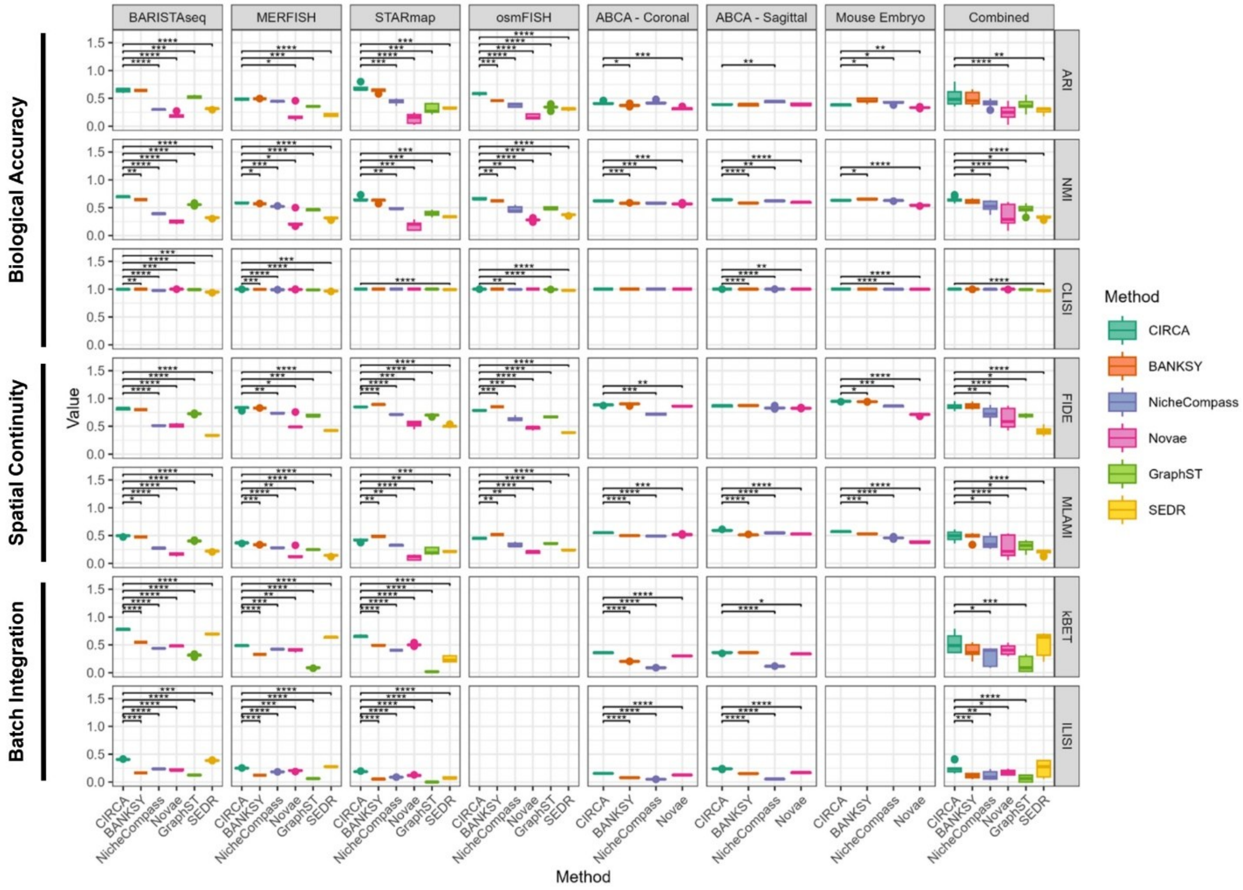


**Supplementary Figure S1:** Benchmarking results for CIRCA and five competing methods across seven annotated datasets on individual metrics measuring biological accuracy, spatial continuity, and multi-slide integration. Boxplots depict performance across 5 random seeds; the center line denotes the median, box limits denote the upper and lower quartiles, and whiskers denote 1.5x the interquartile range; Welch's t-test (within dataset) and mixed effects model (combined) Holm adjusted p-values: * p<0.05, ** p<0.01, *** p<0.001, **** p<0.0001.

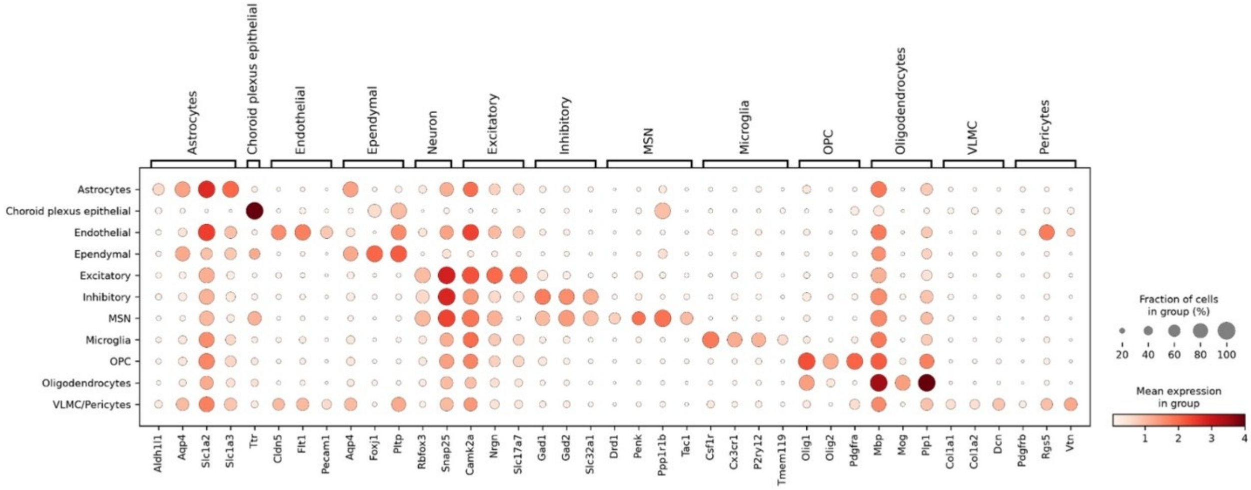


**Supplementary Figure S2:** Mean expression of broad cell type marker genes (indicated by the brackets along the top) across major cell types identified by clustering CIRCA's cell-state embeddings. Abbreviations: MSN, medium spiny neuron; OPC, oligodendrocyte precursor cell; VLMC, vascular leptomeningeal cells.

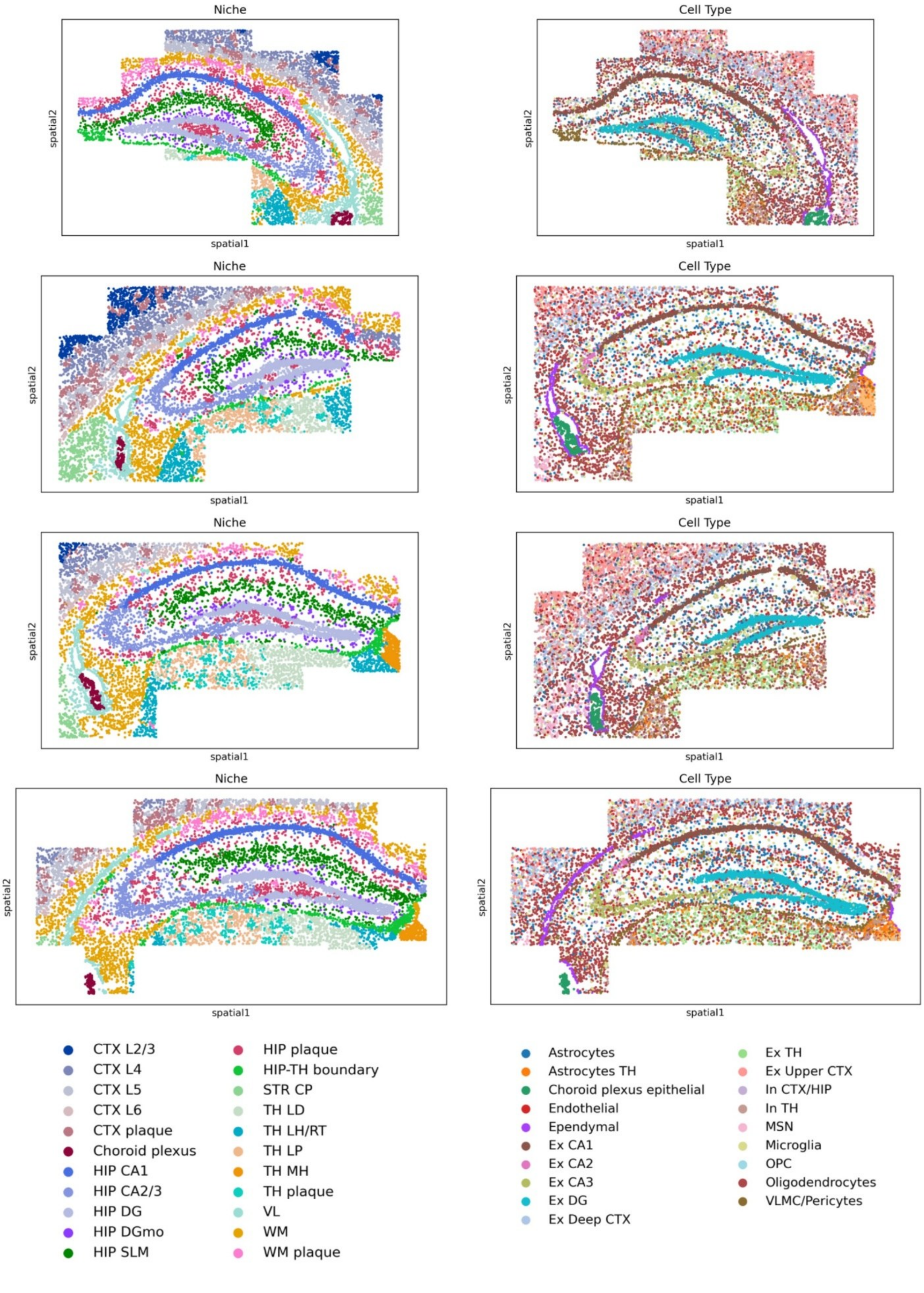
Niche
Cell Type
spatial1
spatial2
CTX L2/3
CTX L4
CTX L5
CTX L6
CTX plaque
Choroid plexus
HIP CA1
HIP CA2/3
HIP DG
HIP DGmo
HIP SLM
HIP plaque
HIP-TH boundary
STR CP
TH LD
TH LH/RT
TH LP
TH MH
TH plaque
VL
WM
WM plaque
Astrocytes
Astrocytes TH
Choroid plexus epithelial
Endothelial
Ependymal
Ex CA1
Ex CA2
Ex CA3
Ex DG
Ex Deep CTX
Ex TH
Ex Upper CTX
In CTX/HIP
In TH
MSN
Microglia
OPC
Oligodendrocytes
VLMC/Pericytes

**Supplementary Figure S3:** Spatial distributions of CIRCA-identified niches (left) and cell types (right) in all hippocampal sections.

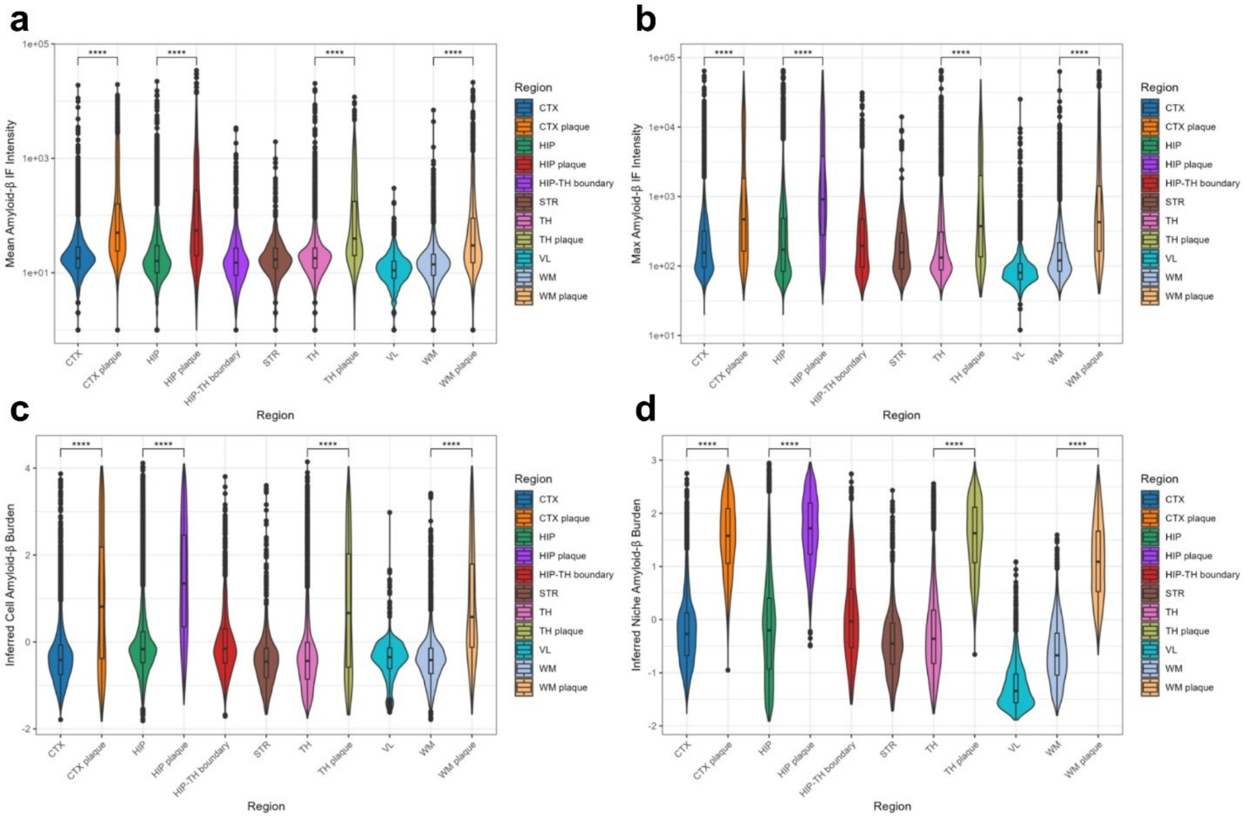


**Supplementary Figure S4:** Violin plots depicting the distribution of Aβ immunofluorescence intensity features (**a,b**) and CIRCA-estimated Aβ latent factors (**c,d**) across brain regions and plaque niches identified by CIRCA. In box plots, the center line denotes the median, box limits denote the upper and lower quartiles, and whiskers denote 1.5x the interquartile range; Wilcoxon rank sum test raw p-values: * p<0.05, ** p<0.01, *** p<0.001, **** p<0.0001. Abbreviations: CTX, cortex; HIP, hippocampus; STR, striatum; TH, thalamus; VL, lateral ventricle; WM, white matter.

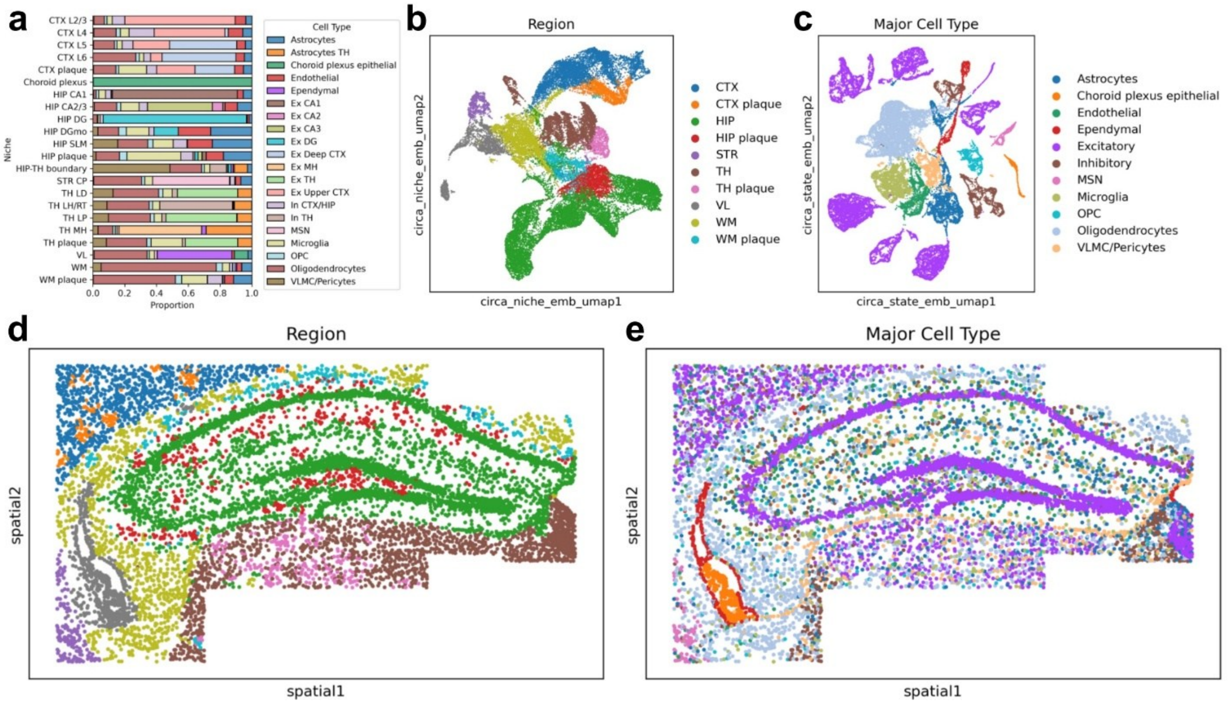


**Supplementary Figure S5: a)** Cell type composition of each niche. **b-c)** UMAPs of CIRCA niche (**b**) and cell-state (**c**) embeddings annotated with broader region (**b**) and major cell type (**c**) labels derived from Leiden clustering. **d-e)** Spatial distributions of CIRCA-identified regions (**d**) and major cell types (**e**) in a representative hippocampal section. Abbreviations: CTX, cortex; HIP, hippocampus; MSN, medium spiny neuron; OPC, oligodendrocyte precursor cell; STR, striatum; TH, thalamus; VL, lateral ventricle; VLMC, vascular leptomeningeal cells; WM, white matter.

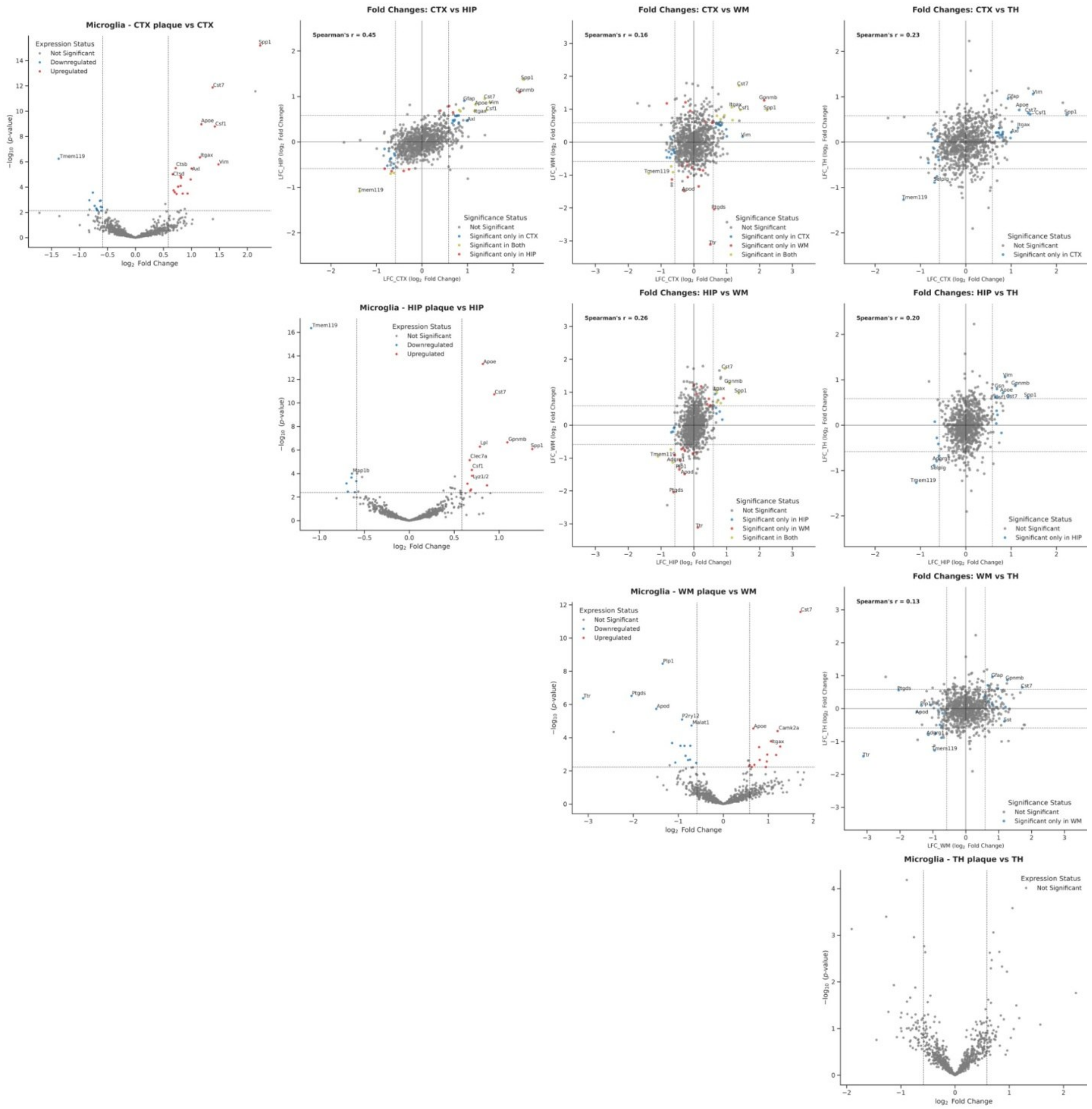

**Supplementary Figure S6:** Pair plots depicting pseudobulk differential expression results comparing plaque-associated microglia to non-plaque microglia in brain regions with plaque-associated niches. A log fold change (LFC) threshold of ±0.5 and FDR-adjusted p-value threshold of 0.05 were used to determine significance. The diagonal displays volcano plots for each comparison, while the off-diagonal scatter plots display the relationship between LFC values for regional comparisons.

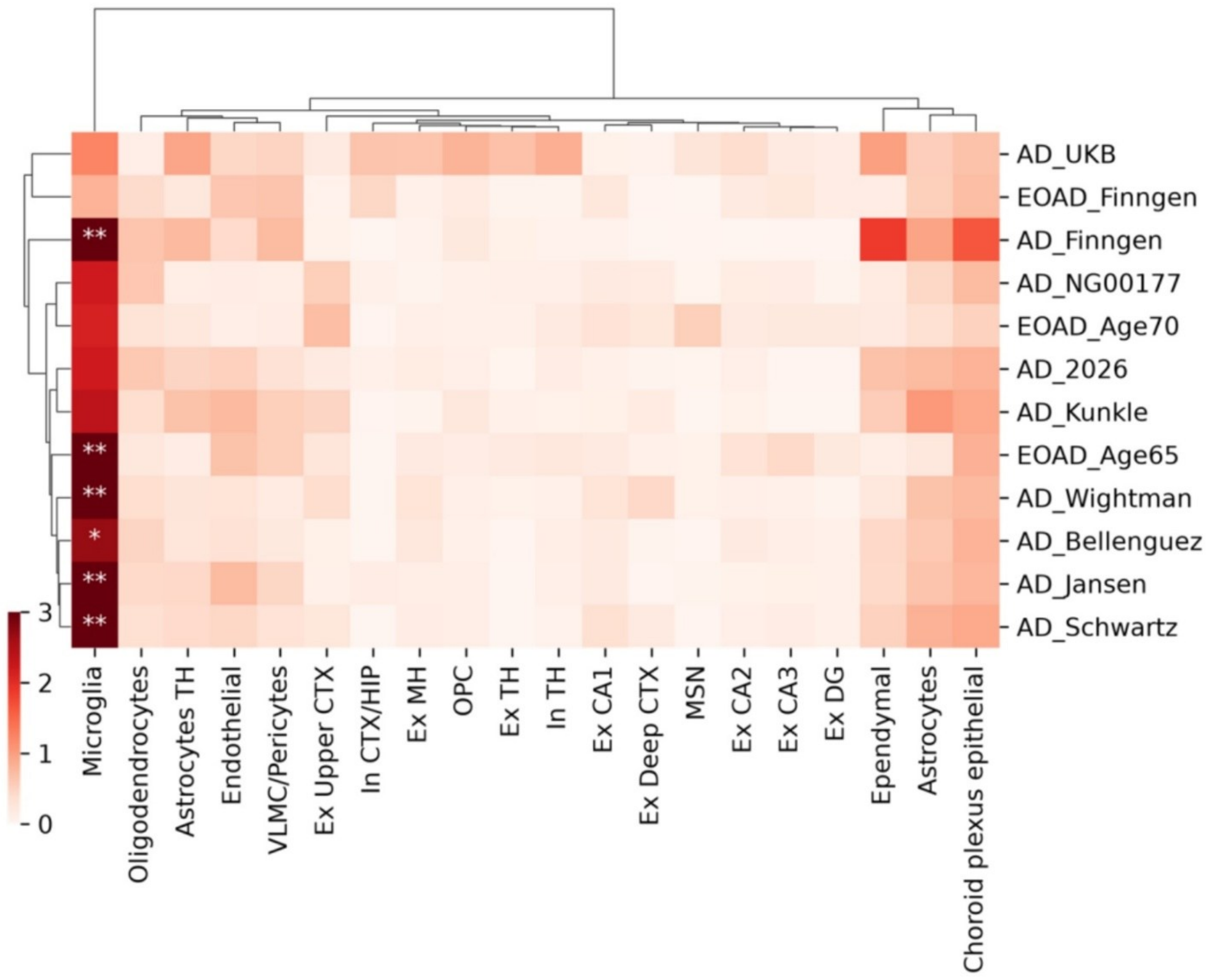


**Supplementary Figure S7:** scDRS enrichment for GWAS-derived AD genetic risk across CIRCA cell types. FDR: * <0.1, ** <0.05.

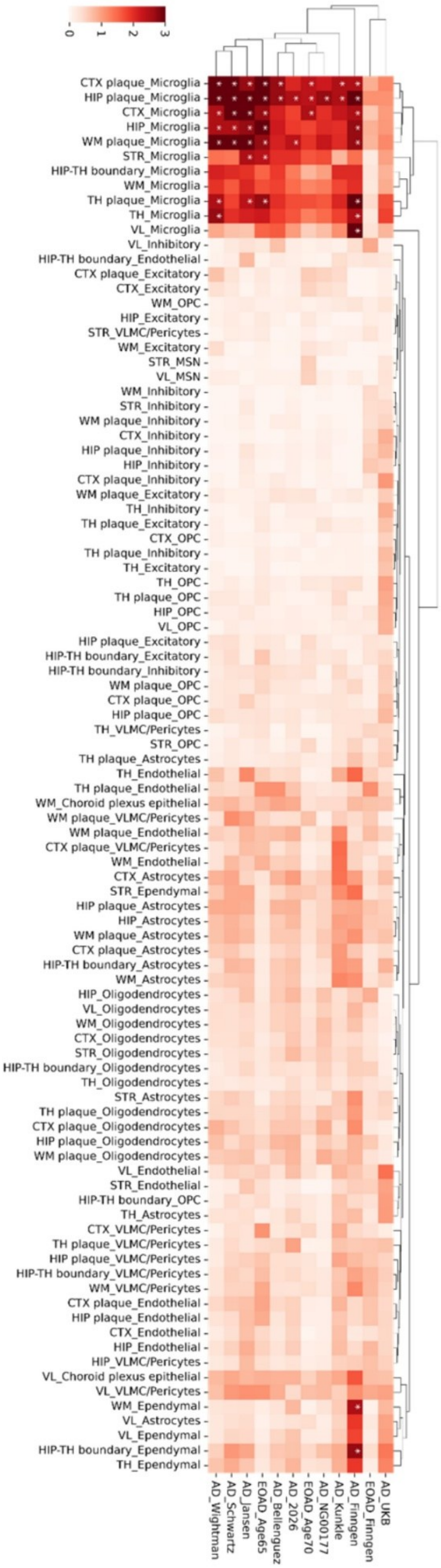

0
1
2
3
CTX plaque_Microglia
HIP plaque_Microglia
CTX_Microglia
HIP_Microglia
WM plaque_Microglia
STR_Microglia
HIP-TH boundary_Microglia
WM_Microglia
TH plaque_Microglia
TH_Microglia
VL_Microglia
VL_Inhibitory
HIP-TH boundary_Endothelial
CTX plaque_Excitatory
CTX_Excitatory
WM_OPC
HIP_Excitatory
STR_VLMC/Pericytes
WM_Excitatory
STR_MSN
VL_MSN
WM_Inhibitory
STR_Inhibitory
WM plaque_Inhibitory
CTX_Inhibitory
HIP plaque_Inhibitory
HIP_Inhibitory
CTX plaque_Inhibitory
WM plaque_Excitatory
TH_Inhibitory
TH plaque_Excitatory
CTX_OPC
TH plaque_Inhibitory
TH_Excitatory
TH_OPC
TH plaque_OPC
HIP_OPC
VL_OPC
HIP plaque_Excitatory
HIP-TH boundary_Excitatory
HIP-TH boundary_Inhibitory
WM plaque_OPC
CTX plaque_OPC
HIP plaque_OPC
TH_VLMC/Pericytes
STR_OPC
TH plaque_Astrocytes
TH_Endothelial
TH plaque_Endothelial
WM_Choroid plexus epithelial
WM plaque_VLMC/Pericytes
WM plaque_Endothelial
CTX plaque_VLMC/Pericytes
WM_Endothelial
CTX_Astrocytes
STR_Ependymal
HIP plaque_Astrocytes
HIP_Astrocytes
WM plaque_Astrocytes
CTX plaque_Astrocytes
HIP-TH boundary_Astrocytes
WM_Astrocytes
HIP_Oligodendrocytes
VL_Oligodendrocytes
WM_Oligodendrocytes
CTX_Oligodendrocytes
STR_Oligodendrocytes
HIP-TH boundary_Oligodendrocytes
TH_Oligodendrocytes
STR_Astrocytes
TH plaque_Oligodendrocytes
CTX plaque_Oligodendrocytes
HIP plaque_Oligodendrocytes
WM plaque_Oligodendrocytes
VL_Endothelial
STR_Endothelial
HIP-TH boundary_OPC
TH_Astrocytes
CTX_VLMC/Pericytes
TH plaque_VLMC/Pericytes
HIP plaque_VLMC/Pericytes
HIP-TH boundary_VLMC/Pericytes
WM_VLMC/Pericytes
CTX plaque_Endothelial
HIP plaque_Endothelial
CTX_Endothelial
HIP_Endothelial
HIP_VLMC/Pericytes
VL_Choroid plexus epithelial
VL_VLMC/Pericytes
WM_Ependymal
VL_Astrocytes
VL_Ependymal
HIP-TH boundary_Ependymal
TH_Ependymal
AD_Wightman
AD_Schwartz
AD_Jansen
EOAD_Age65
AD_Bellenguez
AD_2026
EOAD_Age70
AD_NG00177
AD_Kunkle
AD_Finngen
EOAD_Finngen
AD_UKB

**Supplementary Figure S8:** scDRS enrichment for GWAS-derived AD genetic risk across major cell types stratified by region. FDR: * <0.1, ** <0.05.

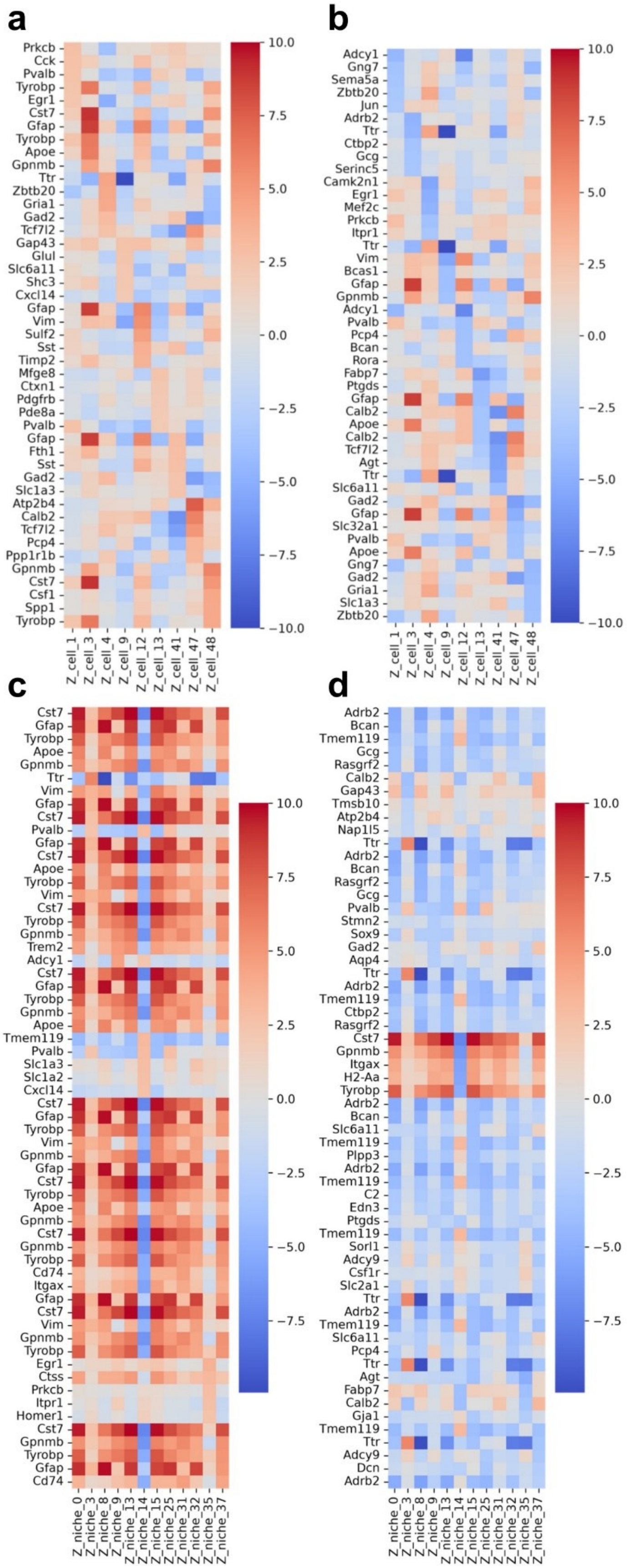
a
Prkcb
Cck
Pvalb
Tyrobp
Egr1
Cst7
Gfap
Tyrobp
Apoe
Gpnmb
Ttr
Zbtb20
Gria1
Gad2
Tcf7l2
Gap43
Glul
Slc6a11
Shc3
Cxcl14
Gfap
Vim
Sulf2
Sst
Timp2
Mfge8
Ctxn1
Pdgfrb
Pde8a
Pvalb
Gfap
Fth1
Sst
Gad2
Slc1a3
Atp2b4
Calb2
Tcf7l2
Pcp4
Ppp1r1b
Gpnmb
Cst7
Csf1
Spp1
Tyrobp
Z_cell_1
Z_cell_3
Z_cell_4
Z_cell_9
Z_cell_12
Z_cell_13
Z_cell_41
Z_cell_47
Z_cell_48
10.0
7.5
5.0
2.5
0.0
−2.5
−5.0
−7.5
−10.0
b
Adcy1
Gng7
Sema5a
Zbtb20
Jun
Adrb2
Ttr
Ctbp2
Gcg
Serinc5
Camk2n1
Egr1
Mef2c
Prkcb
Itpr1
Ttr
Vim
Bcas1
Gfap
Gpnmb
Adcy1
Pvalb
Pcp4
Bcan
Rora
Fabp7
Ptgds
Gfap
Calb2
Apoe
Calb2
Tcf7l2
Agt
Ttr
Slc6a11
Gad2
Gfap
Slc32a1
Pvalb
Apoe
Gng7
Gad2
Gria1
Slc1a3
Zbtb20
Z_cell_1
Z_cell_3
Z_cell_4
Z_cell_9
Z_cell_12
Z_cell_13
Z_cell_41
Z_cell_47
Z_cell_48
10.0
7.5
5.0
2.5
0.0
−2.5
−5.0
−7.5
−10.0
c
Cst7
Gfap
Tyrobp
Apoe
Gpnmb
Ttr
Vim
Gfap
Cst7
Pvalb
Gfap
Cst7
Apoe
Tyrobp
Vim
Cst7
Tyrobp
Gpnmb
Trem2
Adcy1
Cst7
Gfap
Tyrobp
Gpnmb
Apoe
Tmem119
Pvalb
Slc1a3
Slc1a2
Cxcl14
Cst7
Gfap
Tyrobp
Vim
Gpnmb
Gfap
Cst7
Tyrobp
Apoe
Gpnmb
Cst7
Gpnmb
Tyrobp
Cd74
Itgax
Gfap
Cst7
Vim
Gpnmb
Tyrobp
Egr1
Ctss
Prkcb
Itpr1
Homer1
Cst7
Gpnmb
Tyrobp
Gfap
Cd74
Z_niche_0
Z_niche_3
Z_niche_8
Z_niche_9
Z_niche_13
Z_niche_14
Z_niche_15
Z_niche_25
Z_niche_31
Z_niche_32
Z_niche_35
Z_niche_37
10.0
7.5
5.0
2.5
0.0
−2.5
−5.0
−7.5
d
Adrb2
Bcan
Tmem119
Gcg
Rasgrf2
Calb2
Gap43
Tmsb10
Atp2b4
Nap1l5
Ttr
Adrb2
Bcan
Rasgrf2
Gcg
Pvalb
Stmn2
Sox9
Gad2
Aqp4
Ttr
Adrb2
Tmem119
Ctbp2
Rasgrf2
Cst7
Gpnmb
Itgax
H2-Aa
Tyrobp
Adrb2
Bcan
Slc6a11
Tmem119
Plpp3
Adrb2
Tmem119
C2
Edn3
Ptgds
Tmem119
Sorl1
Adcy9
Csf1r
Slc2a1
Ttr
Adrb2
Tmem119
Slc6a11
Pcp4
Ttr
Agt
Fabp7
Calb2
Gja1
Tmem119
Ttr
Adcy9
Dcn
Adrb2
Z_niche_0
Z_niche_3
Z_niche_8
Z_niche_9
Z_niche_13
Z_niche_14
Z_niche_15
Z_niche_25
Z_niche_31
Z_niche_32
Z_niche_35
Z_niche_37
10.0
7.5
5.0
2.5
0.0
−2.5
−5.0
−7.5

**Supplementary Figure S9: a,b)** Heatmap depicting the genes with the top five positive (**a**) and negative (**b**) attributions for each cell-level factor in **Fig. 4c**. **c,d)** Heatmap depicting the genes with the top five positive (**c**) and negative (**d**) attributions for each niche-level factor in **Fig. 4c**.

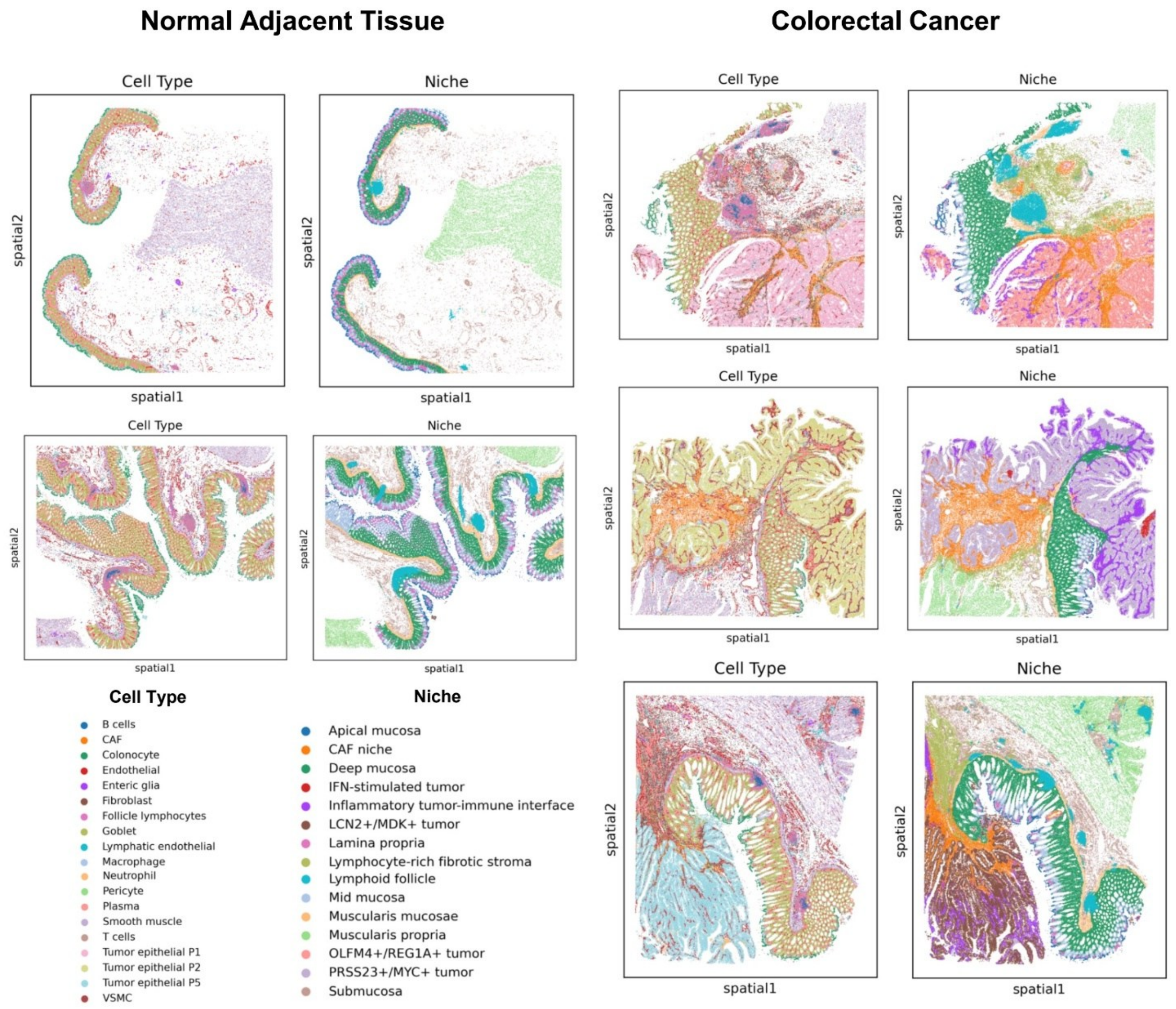


**Supplementary Figure S10:** Spatial distributions of CIRCA-identified niches (right) and cell types (left) in all normal adjacent tissue and colorectal cancer sections.

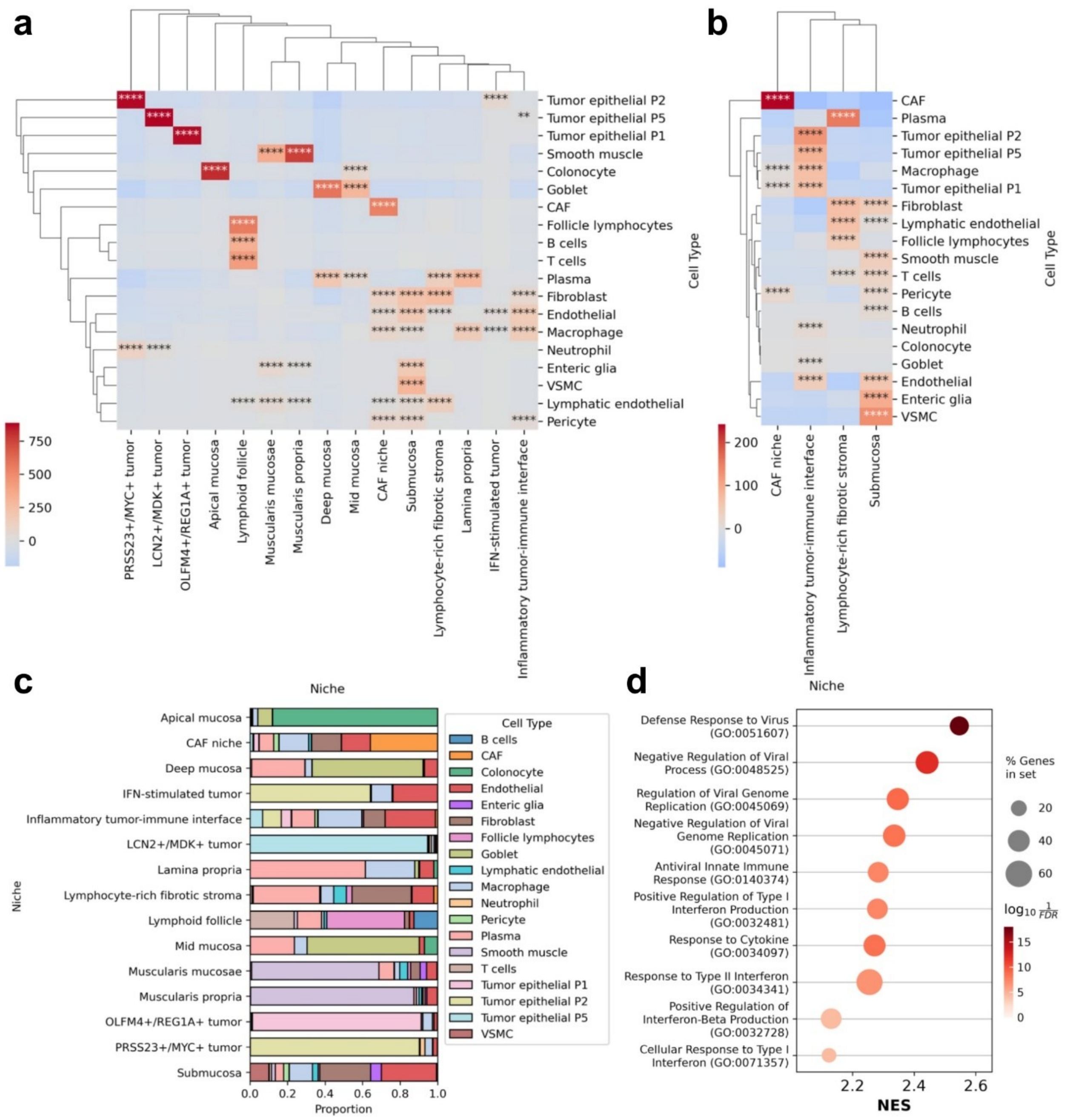


**Supplementary Figure S11: a,b)** Adjusted standardized residuals from chi-square tests of cell type-niche association across all niches (**a**) and across tumor-associated stromal niches and normal submucosa (**b**); Bonferroni-adjusted post hoc p-values: * p<0.05, ** p<0.01, *** p<0.001, **** p<0.0001. **c)** Cell type composition of each niche. Notably, each tumor niche is composed primarily of patient-specific malignant cells. **d)** The top ten GO:BP terms with significant enrichment (FDR $< 0.05$) in the interferon-stimulated tumor niche. Abbreviations: CAF, cancer-associated fibroblast; GO:BP, Gene Ontology: Biological Process; IFN, interferon; VSMC, vascular smooth muscle cells.

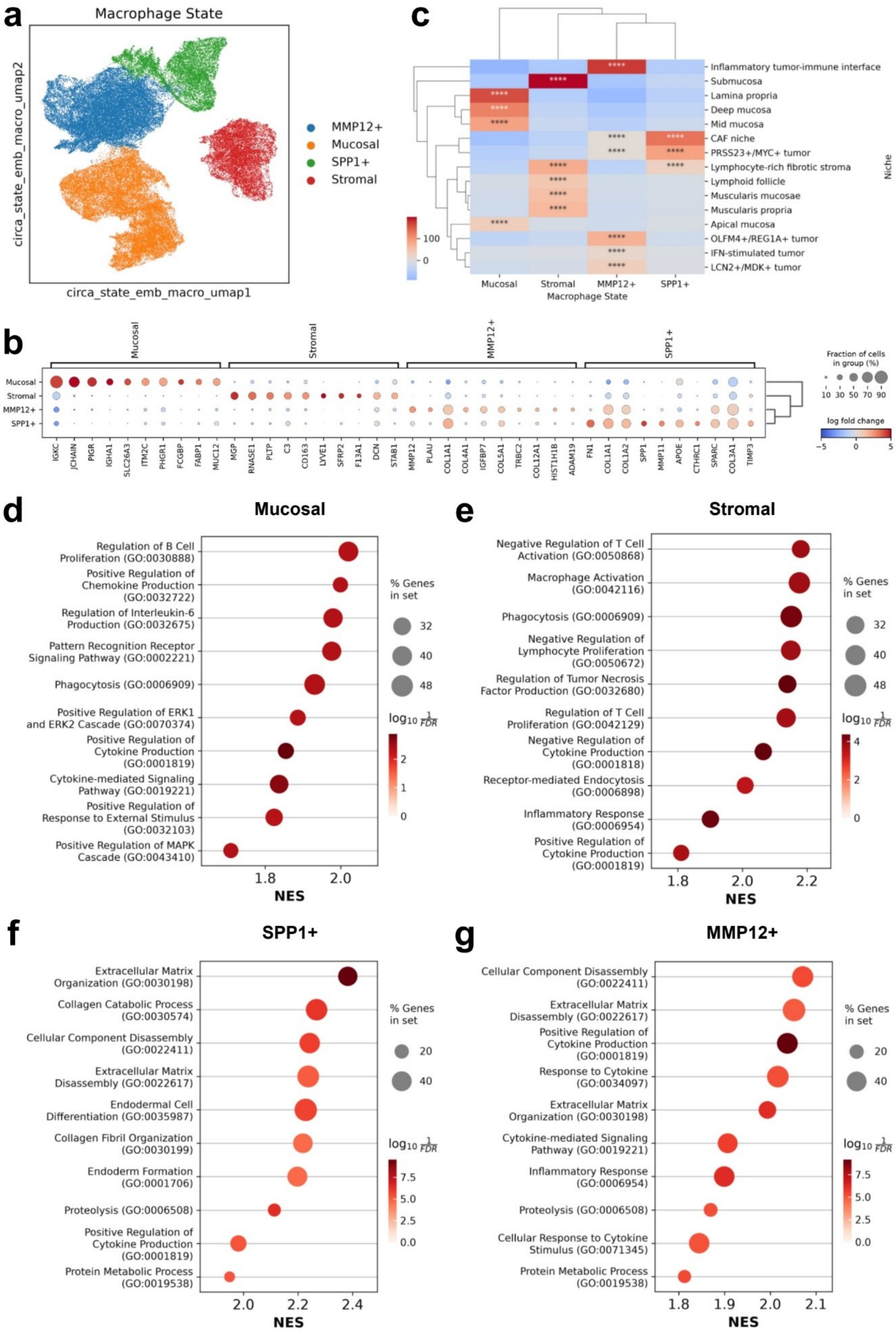
a
Macrophage State
circa_state_emb_macro_umap2
circa_state_emb_macro_umap1
MMP12+
Mucosal
SPP1+
Stromal
c
Inflammatory tumor-immune interface
Submucosa
Lamina propria
Deep mucosa
Mid mucosa
CAF niche
PRSS23+/MYC+ tumor
Lymphocyte-rich fibrotic stroma
Lymphoid follicle
Muscularis mucosae
Muscularis propria
Apical mucosa
OLFM4+/REG1A+ tumor
IFN-stimulated tumor
LCN2+/MDK+ tumor
Niche
100
0
Mucosal
Stromal
MMP12+
SPP1+
Macrophage State
b
Mucosal
Stromal
MMP12+
SPP1+
IGKC
JCHAIN
PIGR
IGHA1
SLC26A3
ITM2C
PHGR1
FCGBP
FABP1
MUC12
MGP
RNASE1
PLTP
C3
CD163
LYVE1
SFRP2
F13A1
DCN
STAB1
MMP12
PLAU
COL1A1
COL4A1
IGFBP7
COL5A1
TRBC2
COL12A1
HIST1H1B
ADAM19
FN1
COL1A1
COL1A2
SPP1
MMP11
APOE
CTHRC1
SPARC
COL3A1
TIMP3
Fraction of cells in group (%)
10 30 50 70 90
log fold change
−5 0 5
d
Mucosal
Regulation of B Cell Proliferation (GO:0030888)
Positive Regulation of Chemokine Production (GO:0032722)
Regulation of Interleukin-6 Production (GO:0032675)
Pattern Recognition Receptor Signaling Pathway (GO:0002221)
Phagocytosis (GO:0006909)
Positive Regulation of ERK1 and ERK2 Cascade (GO:0070374)
Positive Regulation of Cytokine Production (GO:0001819)
Cytokine-mediated Signaling Pathway (GO:0019221)
Positive Regulation of Response to External Stimulus (GO:0032103)
Positive Regulation of MAPK Cascade (GO:0043410)
1.8
2.0
NES
% Genes in set
32
40
48
$\log_{10}\frac{1}{FDR}$
2
1
0
e
Stromal
Negative Regulation of T Cell Activation (GO:0050868)
Macrophage Activation (GO:0042116)
Phagocytosis (GO:0006909)
Negative Regulation of Lymphocyte Proliferation (GO:0050672)
Regulation of Tumor Necrosis Factor Production (GO:0032680)
Regulation of T Cell Proliferation (GO:0042129)
Negative Regulation of Cytokine Production (GO:0001818)
Receptor-mediated Endocytosis (GO:0006898)
Inflammatory Response (GO:0006954)
Positive Regulation of Cytokine Production (GO:0001819)
1.8
2.0
2.2
NES
% Genes in set
32
40
48
$\log_{10}\frac{1}{FDR}$
4
2
0
f
SPP1+
Extracellular Matrix Organization (GO:0030198)
Collagen Catabolic Process (GO:0030574)
Cellular Component Disassembly (GO:0022411)
Extracellular Matrix Disassembly (GO:0022617)
Endodermal Cell Differentiation (GO:0035987)
Collagen Fibril Organization (GO:0030199)
Endoderm Formation (GO:0001706)
Proteolysis (GO:0006508)
Positive Regulation of Cytokine Production (GO:0001819)
Protein Metabolic Process (GO:0019538)
2.0
2.2
2.4
NES
% Genes in set
20
40
$\log_{10}\frac{1}{FDR}$
7.5
5.0
2.5
0.0
g
MMP12+
Cellular Component Disassembly (GO:0022411)
Extracellular Matrix Disassembly (GO:0022617)
Positive Regulation of Cytokine Production (GO:0001819)
Response to Cytokine (GO:0034097)
Extracellular Matrix Organization (GO:0030198)
Cytokine-mediated Signaling Pathway (GO:0019221)
Inflammatory Response (GO:0006954)
Proteolysis (GO:0006508)
Cellular Response to Cytokine Stimulus (GO:0071345)
Protein Metabolic Process (GO:0019538)
1.8
1.9
2.0
2.1
NES
% Genes in set
20
40
$\log_{10}\frac{1}{FDR}$
7.5
5.0
2.5
0.0

**Supplementary Figure S12: a)** UMAP of CIRCA cell-state embeddings for macrophages annotated by macrophage state. **b)** Top ten differentially expressed genes for each macrophage state. **c)** Adjusted standardized residuals from chi-square tests of macrophage state-niche association; Bonferroni-adjusted post hoc p-values: * $p<0.05$, ** $p<0.01$, *** $p<0.001$, **** $p<0.0001$. **d-g)** The top ten GO:BP terms with significant enrichment (FDR < 0.05) in the mucosal (**d**), stromal (**e**), SPP1+ (**f**), and MMP12+ (**g**) macrophage states. Abbreviations: CAF, cancer-associated fibroblast; GO:BP, Gene Ontology: Biological Process; IFN, interferon.
